# Chronic pancreatitis patient-derived organoids reveal new paths to precision therapeutics

**DOI:** 10.1101/2024.10.30.620903

**Authors:** Victoria Osorio-Vasquez, Jonathan Zhu, Jan C. Lumibao, Kathryn Lande, Kristina L. Peck, McKenna K. Stamp, Shira R. Okhovat, Hyemin Song, Satoshi Ogawa, Casie Kubota, Vasiliki Pantazopoulou, Yang Dai, Angelica Rock, Chelsea Bottomley, Ethan Thomas, Jasper Hsu, Araceli Herrera Morales, Alexandra Fowler, T’Onj McGriff, K. Garrett Evensen, Siri Larsen, Muhamad Abdulla, Phil Greer, Jessica Gibson, Michael Downes, Ronald Evans, Andrew M. Lowy, David C. Whitcomb, Jingjing Zou, Alfredo Molinolo, Tae Gyu Oh, Rebekah White, Melena Bellin, Herve Tiriac, Dannielle D. Engle

## Abstract

Chronic pancreatitis (CP) affects ∼3 million people worldwide, yet altering the course of disease is challenging. We developed a patient-derived organoid (PDO) platform to investigate the molecular pathogenesis of this disease and identify therapeutic strategies. We generated 36 PDOs from patients with idiopathic, hereditary, and alcohol-related CP with high genetic concordance. PDOs retained inflammation-associated transcriptional and proteomic features. Transcriptomic profiling revealed three molecular subtypes of CP independent of etiology. We discovered widespread dysfunction of the cystic fibrosis transmembrane conductance regulator (CFTR) in half of the CP PDOs, including those with wildtype CFTR. Clinically available CFTR modulators stabilized mutant or wildtype CFTR, restored CFTR function, and decreased mitogenic and inflammatory signaling. This work provides the first comprehensive PDO platform for modeling CP. We demonstrate the utility of this platform for precision therapeutic investigations. Our findings reveal CFTR modulators as a broadly applicable and effective therapeutic strategy.

## Introduction

Pancreatitis, or pancreatic inflammation, is a continuum of diseases that may progress from acute to chronic stages. More than 300,000 Americans and 3 million individuals globally have chronic pancreatitis (CP)^1–3^. CP is characterized by atrophy, fibrosis, pancreatic exocrine insufficiency, diabetes mellitus, and intractable pain. Current treatment is largely supportive and often fails to modify disease progression, sometimes requiring surgical interventions such as partial or total pancreatectomy for pain management. CP may arise from numerous environmental and genetic factors. It is classified using the Toxic-metabolic, Idiopathic, Genetic, Autoimmune, Recurrent and severe acute pancreatitis and Obstructive (TIGAR-O v2) framework^4^. The most common identifiable causes of CP include tobacco use (46-60%)^5,6^, gallstones (14-37%)^7,8^, alcohol (7-34%)^5,9^, and genetics (10%)^5^, though a striking 28-80% of cases remain idiopathic^10^. Hereditary CP is associated with pathogenic germline mutations in protease serine 1 (*PRSS1*) (68-80%)^11–14^, serine protease inhibitor kazal type 1 (*SPINK1*) (1-8.8%)^10–13^, cystic fibrosis transmembrane conductance regulator (*CFTR*) (16.5%)^13^, and other genes.

Pancreatitis initiation can be traced to dysfunction in both acinar and ductal cells. *PRSS1* and *SPINK1* mutations lead to hyperactivation of digestive enzymes, leading to acinar cell injury and inflammation^15–20^. Mutations in *CFTR* disrupt bicarbonate secretion in the pancreas. Ductal cells normally secrete 1-2L of 140mM sodium bicarbonate per day^21^, which is essential for mucin production, flushing digestive enzymes into the duodenum, preventing premature digestive enzyme activation, and neutralizing acidic chyme from the stomach^21–23^. Unlike cystic fibrosis, CFTR variants associated with hereditary CP are often heterozygous^24^. Up to 37% of hereditary CP patients harbor *CFTR* mutations^25–32^. Importantly, 50% of idiopathic CP patients harbor *CFTR* alterations when extensive genotyping is performed^21,25–34^. Many *CFTR* variants associated with hereditary CP selectively impair bicarbonate, but not chloride transductance. Altogether, CFTR dysfunction in the pancreas, distinct from classical cystic fibrosis, may underlie a significant proportion of CP and represents an underexplored therapeutic target.

With more than 2,000 mutations annotated for *CFTR*, the response of most *CFTR* variants to modulators are unknown. Of those with response information, this testing has been primarily performed for cystic fibrosis-related variants in non-pancreas relevant contexts. Further, cystic fibrosis patient responses to CFTR therapies are highly variable due to undefined mechanisms^35–40^. Limited information is available regarding CFTR modulator response in the context of hereditary pancreatitis associated *CFTR* mutations. Delineating the genotypic and phenotypic relationships of pancreatitis-associated CFTR alterations with modulator response is essential. But these investigations have been hampered by the limited availability of renewable human models. Unfortunately, despite sharing 78% homology, mouse CFTR does not respond to human CFTR modulators, complicating mechanistic investigations in mouse models^41^. Further, mouse bicarbonate secretion is notably different than humans, reaching only 50mM in pancreatic secretions^37^, limiting our ability to make conclusions relevant to human physiology.

Three-dimensional patient-derived organoids (PDOs) have emerged as transformative models for studying pancreas biology. PDOs can be generated from normal human pancreas and pancreatic ductal adenocarcinoma with high efficiency ^42,43^, far surpassing earlier monolayer or iPSC-based platforms^44^. PDOs are expandable, cryopreservable, and genetically stable, enabling in-depth molecular and functional analyses^43,45–50^. In pancreatic ductal adenocarcinoma (PDA), PDOs faithfully recapitulate transcriptomic and genetic subtypes, predict treatment response, and inform mechanisms of resistance^43,45–50^. Despite lacking a microenvironment, PDOs retain remarkable predictive power due to the central role of epithelial-intrinsic signaling in disease behavior^51,52^. PDO models of CP remain rare. A pediatric cohort (n=9) has been described^53^, but generation efficiency, molecular fidelity, and translational potential remain unknown. To address this gap, we created a comprehensive biobank of CP PDOs encompassing all major etiologies and benchmarked their ability to model disease features and guide treatment strategies.

We report a high PDO generation success rate (70.5%, *n*=36) and strong concordance with matched primary tissues at the DNA, RNA, and protein levels. Genetic analyses of 24 PDOs identified germline pathogenic variants in 2 out of 8 idiopathic cases. In addition, rare somatic variant analyses identified 2 CP PDOs with alterations associated with pancreatic ductal adenocarcinoma, including *KRAS* and *TP53*. These oncogenic alterations were not detectable in primary samples in which low cellularity limits the ability to detect somatic mutations in the epithelial compartment. CP PDOs exhibit elevated fibroinflammatory gene and protein expression profiles despite being removed from the pancreatic microenvironment, suggesting that ductal epithelial cell intrinsic features contribute to inflammation. Strikingly, half of the PDOs tested displayed CFTR functional impairment despite 2 out of the 4 dysfunctional lines harboring wildtype CFTR. Treatment with clinically available CFTR modulators restored ductal function and suppressed pro-inflammatory and mitogenic signaling, suggesting that CFTR modulation may benefit a broad segment of CP patients. Together, this study presents a robust and scalable platform for dissecting CP pathophysiology and accelerating precision medicine efforts.

## Methods

### Human Specimens

Normal pancreata were obtained from the Lifesharing organ donation program. The average age of the hNP PDOs is 56 years of age. Pancreatitis tissues were obtained from the University of Minnesota (UMN) and University of California San Diego (UCSD) Biorepository and Tissue Technology Shared Resource (BTTSR). De-identified pancreatitis tissues were obtained from the University of Minnesota (UMN) or the UCSD BTTSR with approval from their respective Institutional Review Boards. Written and informed consent or parental assent, as age appropriate, was obtained prior to acquisition of tissue from all patients. Age and/or age ranges will not be provided for pediatric patients as the rarity of the procedure in this patient population could itself be identifying. Specimens were obtained from patients with chronic pancreatitis at UCSD undergoing either partial pancreatectomy (distal pancreatectomy or pancreaticoduodenectomy) or pancreatic head debridement during pancreatic drainage procedures for symptomatic relief. Excess tissue was collected fresh into cold media for organoid generation. When available, pancreatic juice was collected in parallel by needle aspiration of pancreatic duct during drainage procedures. Specimens were obtained from patients undergoing total pancreatectomy and islet auto-transplantation at UMN. During pancreatectomy, the blood supply was preserved to the pancreas as long as possible to avoid prolonged ischemia. The entire pancreas, spleen, and duodenum were removed, and the gastrointestinal and biliary continuity is restored by a Roux-en-Y approach. Islet isolation was performed by infusing the pancreas with collagenase and neutral protease for enzymatic digestion, followed by mechanical disruption using the semi-automated method of Ricordi. The islets were then transported back to the operating room where they were infused via the portal vein on the same day as the pancreatectomy procedure. Residual exocrine pancreas tissue following islet isolation was collected for tissue research. Tissue specimens were catalogued, imaged, and processed (i.e. histology, organoid generation, flash frozen, and viably cryopreserved).

### Histology and immunohistochemistry

Tissues were fixed in 10% neutral buffered formalin for 48 hours. Tissues were embedded in paraffin and stained with hematoxylin and eosin or Masson’s trichrome by the Tissue Technology Shared Resource at UCSD supported by the National Cancer Institute Cancer Center Support Grant (CCSG Grant P30CA23100). The following primary antibodies were used for immunohistochemical staining Ki-67 (8D5, Cell Signaling Technologies# 9449), CK19 (Troma-III, Developmental studies hybridoma bank), CFTR (M3A7, Thermo Fischer #MA5-11768), Cancer Antigen 19-9 (MyBioSource #MBS850562). Secondary antibodies used were ImmPRESS® HRP Horse Anti-Mouse IgG Polymer Detection Kit (MP-7402) or ImmPRESS® HRP Goat Anti-Rat IgG Polymer Detection Kit (MP-7404) with ImmPACT® DAB Substrate Kit (HRP) (SK-4105). Staining intensity was semi quantitatively scored as: low, medium, or high using QuPath 0.5.0^54^ and ranking percent positive staining. Slides were imaged using the Olympus VS-120 virtual slide scanning microscope, Olympus BX43 Microscope, or Zeiss Axioscan 7. Zen lite or CellSens was used to visualize and annotate images.

### Pathologist Review of Patient Primary Tissue Specimen

Primary tissue specimen stained with hematoxylin and eosin or Masson’s trichrome were scanned (Aperio-Leica AT2 Scanner, 40x) analyzed and imaged. Severity of fibrosis, acinar atrophy, inflammation and ductal changes were categorized as mild, moderate, or severe as denoted by dilation of ducts, expansion of fibroblast compartment, collagen deposition, and infiltration of immune cells^55^. Alterations in histology due to autolytic changes or signs of cautery from an electro-scalpel.

### Organoid generation and culture

Prior to processing, visible fat, mesentery, and calcifications were removed. Patient specimens were minced and aliquoted for viable tissue cryopreservation for potential regeneration of PDOs, flash frozen for primary tissue DNA extraction, and further dissociation for PDO generation. For organoid generation, tissue was minced and incubated in digestion media (3000 U/mL Collagenase XI + 1000 U/mL hyaluronidase-Stem Cell #07919 in DMEM/F12 diluted 1:4 in Advanced DMEM (Thermo Fischer #12634010) with 1% (v/v) glutamax (Thermo Fischer #35050061), 1% (v/v) penicillin/streptomycin (Life Technologies #15140122), and 1%(v/v) HEPES (1M, pH 7.5)) at 35°C while rotating. If cell lysis became apparent, DNAse I (10 µg/mL, Roche #1128493200) was added. Fractions were collected every 20 minutes for up to 1 hour by allowing large tissue pieces to sediment before supernatant was collected. Dissociation of the remaining large tissue fragments continued by replacing the digestion media. Supernatant was centrifuged at 300x RCF at 4°C for 5 minutes. The first fraction cell pellet collected was resuspended in 1 mL of HBSS (Life Technologies #14175095) with 2% FBS and placed on ice while collecting additional fractions from the digested tissue. After 3 fractions were collected, the supernatant fractions were washed with Advanced DMEM with 1% (v/v) glutamax, 1% (v/v) penicillin/ streptomycin, and 1% (v/v) HEPES prior to resuspension in Matrigel for plating. If larger tissue pieces remained at the end of dissociation, they were either viably cryopreserved (CrySOfree-Sigma-Aldrich #C9249) or plated in Matrigel (Corning #356231) in separate wells. Cells were plated in Corning Matrigel from lots 28323001, 2011001, 0287001, and 1062004 during growth, expansion and experiments. Following Matrigel solidification, complete feeding media was applied to each well.

Murine normal pancreas organoids were cultured in medium previously described^43,50^. Human normal pancreas and human pancreatitis were cultured in human complete feeding medium. Human complete feeding medium was prepared using Advanced DMEM/F12 containing HEPES (1 % (v/v), 1M, pH 7.5), glutamax 1 % (v/v), B27 supplement (1x,Life Technologies #A3582801), and 1% (v/v) penicillin/streptomycin; N-acetylcysteine (1.25 mM, Sigma-Aldrich #A9165), nicotinamide (10 mM, Thermo Fischer #125271000); hEGF (50 ng/mL, R&D Systems #236-EG), FGF-10 (100 ng/mL, R&D Systems #345-FG), gastrin (10 nM, R&D #3006), and A83-01 (500 nM, Tocris Bioscience #2939); rNoggin (100 ug/mL, Qkine LTD #Qk034) and rRSPO1 (10nM Qkine LTD # Qk006); Wnt surrogate (300 pM, ImmunoPrecise Antibodies #N001); PGE2 (1 µM, Tocris Bioscience #2296) and Y-27632 dihydrocholoride (10.5 µM, Tocris Bioscience #1254). Mouse organoid media lacked Wnt3a, PGE2, and Y-27632 and utilized 10% RSPO1 conditioned media and 10% Noggin conditioned media. Organoid nomenclature is defined as: hNP, human normal pancreas; hCP, human chronic pancreatitis, mNP, mouse normal pancreas. All organoid models were routinely tested for *mycoplasma* at Salk Institute.

### Whole mount immunofluorescence and EdU incorporation

Organoids were cultured as described above in µ-slide (Ibidi #80826). Reagents for Click-iT Plus EdU Cell Proliferation Kit for Imaging (Thermo Fischer #C10639) were used in accordance with manufacturer’s protocol. Organoids were treated with EdU for 25 hours by removing half of the media and adding EdU containing media (20 µM). Matrigel domes containing organoids were washed using PBS prior to fixing with 4% PFA (v/v) at 37°C for 10 minutes. After fixation rinsed with excess 3% BSA and removed wash solution. Permeabilized with 0.5% Triton X-100 in PBS for 10 minutes at room temperature. Steps subsequent to Click-it reaction cocktail addition were protected from light. Remaining organoids were washed with 3% BSA in PBS and blocked with IF buffer (0.1% Bovine serum albumin, 0.2% goat serum, 0.2% triton x-100, 0.05% Tween 20:sterile filtered + 10% Goat serum in PBS) for 1 hour. Organoids were incubated with a dilution of 1:200 for CK-19 (Invitrogen #MA5-12663) overnight at room temperature. Organoids were washed for 1 hour prior to incubation with secondary antibody (Jackson ImmunoResearch Laboratories Inc. #115-545-003) and Hoechst for 2 hours at room temperature. Washed organoids for at least 1 hour with IF buffer prior to mounting using Prolong Glass Antifade Mountant (Thermo Fisher P36980) and dried overnight. Imaged using the Yokogawa CQ3000 at 20x. Nuclei and EdU positive nuclei were quantified using Cell Profiler (3.8.1)^56^.

### Immunoblot

Standard techniques were employed for immunoblotting (IB) of human organoids. Organoids were isolated from Matrigel using cell recovery solution (10 mM HEPES 7.5 pH, 1x glutamax (v/v), 12.5 mM EDTA, 12.5 mM EGTA, 4.5 g/L dextrose in PBS w/o Ca and Mg) on ice with 0.5x protease and phosphatase inhibitors (Thermo Fischer # PIA32961). After incubation on ice for 20 minutes, organoids were centrifuged at 0.8 x 1000 RCF at 4°C to pellet organoids and remove cell recovery solution. Cell pellet was resuspended in TNET (1% (v/v) Triton X100, 150 mM NaCl, 5 mM EDTA, 50 mM Tris pH 7.5) with 1.0X protease and phosphatase inhibitors (Life Technologies #A32961) and incubated on ice for 10 minutes before clarifying lysates for whole cell lysate isolation. Protein quantification was carried out using BCA assay (Thermo Scientific #23227). Protein lysates used to probe for CFTR were warmed to 37°C for 1 hour to prevent insolubility, all other blots were performed using lysates heated to 95°C for 5 minutes. Protein lysates were separated on a 4%-12% Bis-Tris NuPage gel (Invitrogen). Western blots were probed with at a dilution of 1:1000 for CFTR (D6W6L, Cell Signaling Technologies #78335), CA19-9 (MyBioSource #MBS850562), pEGFR (Y1068, Cell Signaling Technologies #3777), EGFR (D38B1, Cell Signaling Technologies #4267), pSTAT3 (Y705, Cell Signaling Technologies #D4XC3), STAT3 (D3Z2G, Cell Signaling Technologies #63585), pAKT (S473, Cell Signaling Technologies #4060), AKT (11 E 7,Cell Signaling Technologies #4685), pERK (T202/Y204,Cell Signaling Technologies #4370S), ERK2 (137F5, Cell Signaling Technologies #4695), CA2 (EPR5195,Abcam #ab124687), SPINK1 (CGQG032302A R&D #AF7496), and Cofilin (D3F9, Cell Signaling Technologies #5175). Secondary antibodies used were Peroxidase AffiniPure™ Goat Anti-Mouse IgG (H+L) (Jackson ImmunoResearch #115-035-003), Peroxidase AffiniPure™ Donkey Anti-Rabbit IgG (H+L) (Jackson ImmunoResearch #711-035-152), Peroxidase AffiniPure™ Goat Anti-Rabbit IgG (H+L) (Jackson ImmunoResearch #111-035-003), Sheep IgG Horseradish Peroxidase-conjugated Antibody (R&D Systems #HAF016), IRDye 800 CW Goat Anti-Rabbit IgG (Licor Bio #926-32211) and IRDye 680 CW Goat Anti-Mouse IgG (Licor Bio #926-68070).

hCP35 and hCP44 organoids at 50-70% confluency were treated with Ivacaftor (3 µM-MedChemExpress #HY-13017), combination of Ivacaftor, Elexacaftor (3 µM-MedChemExpress #HY 111772), and Tezacaftor (3 µM-Selleck Chemicals #S7059), or vehicle control for 48 hours in human complete feeding media prior to whole cell lysate collection, as described above.

Western blots were developed using either a film processor (Mini-Med 90-AFP Manufacturing Corporation) or Odyssey M (LiCor). Quantification of immunoblots was conducted using either ImageJ or Empiria Studio.

### WGS library preparation, sequencing, variant calling, and primary tissue-organoid SNP concordance

DNA was isolated from PDOs after at least 3 passages. Isolation of primary tissue DNA was conducted from flash frozen or viably cryopreserved samples. Genomic DNA was isolated from organoids or primary tissue using DNeasy Blood and Tissue Kit (Qiagen #69504). WGS libraries were prepared using Nextera XT (Illumina DNA library) in accordance with the manufacturer’s instructions. Final libraries were evaluated using TapeStation (Agilent). Libraries were sequenced on an Illumina NovaSeq X sequencer for 30 x coverage for primary tissues and PDO.

Raw data was aligned to the hg38 reference genome with bwa mem v0.7.12-r1039. PCR duplicates were marked with Picard v1.130, and germline variants were called with GATK’s haplotypecaller, v4.0.8.1. SNPs and INDELs were filtered with vcftools v0.1.13 the parameters: max-missing=0.5, mac=2, min-meanDP=15, minQ=32. For the purposes of concordance testing, multi-allelic sites and sites with missing genotype calls were discarded.

Concordance was determined using R ‘irr’ package (version 0.84.1) using intraclass correlation coefficient (ICC) using a two-way model with a 0.95 confidence level. As a control, comparison of PDOs from two different patients and found a match rate of 72% with an ICC of 0.4. When comparing a PDO from one patient to the primary tissue of another, the match rate is 74% with an ICC of 0.43.

Germline pathogenic variants were investigated in: ABCG8, APOA5, APOB, CASR, CFTR, CLDN2, CTRC, GGT1, HLA-DQA1, HLA-DQB1, LPL, PRSS1, PRSS1-PRSS2, SLC10A2, SLC26A9 and SPINK1

### RNA isolation and RNA-seq library construction

For RNA-seq experiment, organoids in Matrigel were lysed directly with 1 mL of TRIzol reagent (Thermo Fischer #15596018), and total RNA was extracted using RNeasy Mini Kit (Qiagen #74104). RNA quality control was performed on all samples using Qubit Fluorometer (Invitrogen) and TapeStation (Agilent) before RNA-seq analyses. A total of 200 ng of high-quality RNA (RIN > 8, assessed by Agilent TapeStation) was used for library construction. mRNA was enriched using poly(A) selection, followed by fragmentation. First-and second-strand cDNA synthesis was performed, and 3′ adenylation was carried out prior to ligation of Illumina-compatible adapters containing unique dual barcodes. Adapter-ligated fragments were enriched by limited-cycle PCR amplification. Final libraries were assessed for quality and fragment size distribution using the Agilent Bioanalyzer DNA 1000 kit, and equal amounts of libraries were pooled for sequencing on an Illumina platform. Novaseq 6000 sp reagent kit v 1.5 was used to sequence samples for 300 cycles for 150 bp paired end reads using an Illumina Novoseq 6000.

### snRNA-seq analysis, RNA-seq analysis, PCA visualization, and differential gene expression analysis

Raw reads were quality checked with FASTQC (v0.12.1)^57^. Poor quality 3’ ends and sequencing adapters were trimmed with Cutadapt (v4.9)^58^ and reads shorter than 20nt were filtered. Trimmed reads were again checked with FASTQC, then aligned to human reference genome (hg38 from UCSC with gene annotation gencode.v45.basic.annotation.gtf)^59,60^ and genes quantified with STAR^61^ (2.7.11b). Subsequent analysis was performed in R (version shown below, R Core Team 2024).

The column with 2^nd^ read strand aligned with RNA from the STAR generated expression tables were used since the library prep was strand specific. Expressions were normalized with DESeq2^62^ and the top 500 most variable genes after variance stabilization were used for the PCA plot and heatmap. DESeq2 was also used for differential gene analysis, with cutoffs padj <0.05 and |log_2_FoldChange| >1. These values plotted in the volcano plot for every gene.

Nonnegative matrix factorization exclusively for hCP was performed with the NMF library^63,64^. Ranks 2 through 7 were tested, each with 500 runs to determine the optimal rank. Based on consensus throughout 500 runs, as well as the rank survey scores from the best run each rank, rank 3 was picked as the optimal rank. The consensus shows the best run from the 500 and which samples clustered together but basis was the parameter used to determine the most stable subtypes. Differential gene analysis was performed again, this time separated by basis found with NMF.

Normalized counts from DESeq2 were analyzed with GSEA comparing either hCP with hNAT and hNP, or between hCP bases to determine enriched gene sets^65,66^. GSEA was also used to find gene markers specific to ductal or acinar samples from BioProject PRJNA924346^67^. Each individual PDO transcriptome profile was projected onto the set of ductal or acinar gene markers by single sample GSEA to assess their similarity^68^.

Tumor PDO sample transcriptomes were obtained from NCBI dbGAP accession phs001611.v1.p1^43^. Differentially expressed genes with padj < 0.05 from tumor (hT) vs. normal, and hCP vs hNP and hNAT were integrated to find concordant or discordant expression patterns.

Heatmaps are generated with pheatmap or Graphpad Prism, all other figures are generated with ggplot2. R version and library versions are below.

EGAC00001001732 snRNA-seq data was analyzed^69^, and the ductal cell cluster was interrogated. CP ductal cells contained 903 nuclei from 1 male and 1 female CP patient with unknown etiology, who are 71 and 52 years of age, respectively. NP ductal cells contained 4,432 nuclei from 3 NAT patients (2 female, 1 male, 46, 59, and 77 years of age respectively).

### Fluorescent in situ hybridization

Sections were formalin fix and paraffin embedded. Reagents for the RNAscope Multiplex Fluorescent Reagent kit V2 (#323280) along with RNAscope 4-plex Ancillary Kit for Multiplex Fluorescent Reagent kit V2 (#323120) from Advanced Cell Diagnostics (Newark, CA, USA) were used in accordance with the manufacturer’s instructions using the HybEZ Hybridization system (#241000). Probes were all designed by Advanced Cell Diagnostics: *Gpx3* (#470591-C3), *Aqp1* (#421831-C2), *Bpifb1* (#576511-C4). We used the manual target retrieval method described in appendix B of the protocol for 15 minutes with protease plus treatment for 15 minutes. Fluorescent dyes used TSA Vivid fluorophore 520 (323271-1:3000), 570 (323272-1:3000), 650 (323273-1:3000), and Opal 780 (# FP1501001KT-Akoya Biosciences; Opal TSA-DIG 1:3000; Opal Polaris 780 1:750). Sections were counterstained with DAPI, sealed with a coverslip using ProLong Gold Antifade reagent (Invitrogen #P36930), and then dried overnight covered from light. Imaged using either Fluoview FV 3000 confocal microscope at 30x or whole slide Zeiss Axioscan 7 at 20x.

### Forskolin induced swelling assay

Organoids were seeded into 96-well clear bottom plates in 8 µL domes at a confluency between 50-70%. Organoids were fed human complete medium after seeding. 18-12 hours prior to treatments, media was changed and PGE_2_ was removed from media. For dependency studies, organoids were treated with CFTR and PKA inhibitors for 2 hours prior to forskolin administration (10 µM CFTRinh-172-Selleck Chemicals #S7139; 5 µM PPQ-102-Selleck Chemicals #S1565; 20 µM H-89-Fisher #501873404; 20 µM forskolin-Selleck Chemicals #S2449). Restoration of CFTR function was investigated using CFTR potentiators (10 µM Icenticaftor (QBW251)-Selleck Chemicals #S9973; 10 µM Ivacaftor-MedChemExpress #HY-13017; 10 µM Ivacaftor-Selleck Chemicals #S1144), CFTR corrector (10 µM Lumacaftor-Selleck Chemicals #S1565; 10 µM Elexacaftor-MedChemExpress #HY 111772, Selleck Chemicals #S8851; 10 µM Tezacaftor-Selleck Chemicals #S7059). Mouse normal organoids treated with 20 µM forskolin were used as a positive control for assay. All compounds were dispensed using a Tecan HP D300e in DMSO. Organoids were imaged using brightfield imaging on a Tecan Spark Cyto. Images were taken at time of treatment and 6 hours post treatment with CFTR correctors, potentiators, or forskolin. The area of 10 organoids was measured at these two time points using either QuPath 0.5.0 or ImageJ. The relative area at 6 hours was used for comparison across treatment groups. T-tests and ANOVA statistical tests were conducted.

### Data visualization

Data was visualized using R studio (2024.04.2+764) and GraphPad Prism (9 & 10). Figures were arranged in Adobe Illustrator.

### Data Availability

Raw data for whole genome sequencing and RNA-seq will be publicly available on NCBI the database of genotype and phenotypes (dbGAP) upon final publication of manuscript.

## Results

### Generation of organoids from patients with human chronic pancreatitis (CP) with high efficiency

We received 51 CP surgical specimens (**Table 1, S1, Figures 1A-B, S1A-C, S2A-B**). The patient cohort was predominantly female (65%), with a mean age of 38 years. The mean age was 45 years for patients with idiopathic CP, 25 years for genetic CP, and 47 years for alcohol-related CP. Most patients self-identified as white and non-Hispanic. Idiopathic CP was the most common diagnosis, accounting for 41% of the cohort (**Figure 1B**). Among the 51 patient specimens received, 24 were from patients who underwent genetic testing. Pathogenic variants were reported in CFTR (20%, n=10), PRSS1 (14%, n=7), and SPINK1 (12%, n=6) (**Figure 1C**).

**Figure 1.**
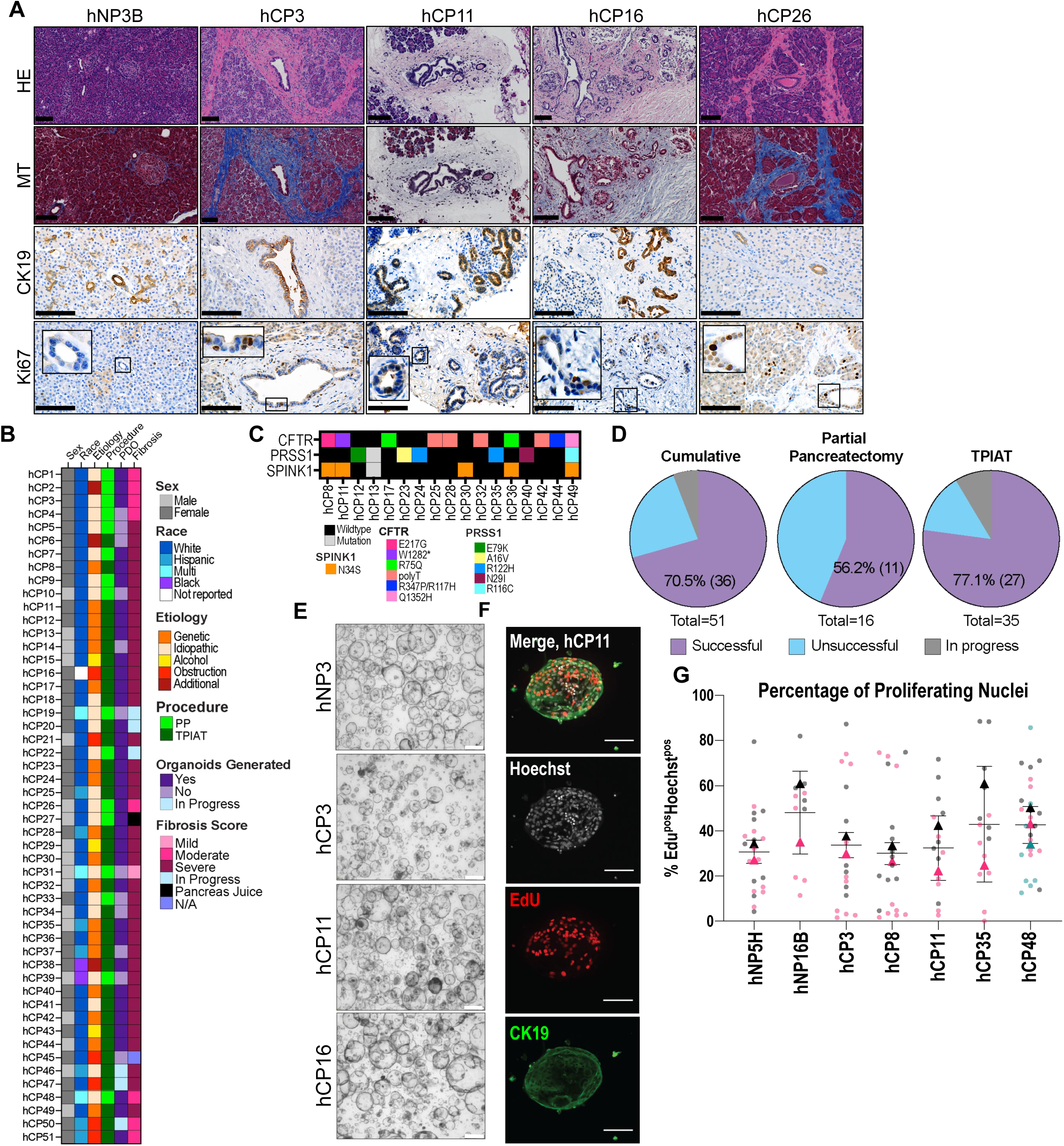
Generation of organoids from patients with human chronic pancreatitis (CP) with high effciency. (A) Representative human normal pancreas (hNP) section displays characteristic ductal, acinar, and islet cells. Masson’s trichrome (MT) staining highlights low degree of fibrosis in NP sample and varying severity in human CP (hCP) samples in blue. NP sample has high levels of cytokeratin 19 staining (CK19) in ductal epithelial cells, while CP samples have an altered ductal cell compartment. Ki-67 expression is low in NP sample and is altered or completely lost in hCP samples. Scale bar 100 µm. For diagram outlining generation procedure and histological features of samples that did not produce organoids see Figure S1. (B) Patient sample cohort demographic, etiology, procedure for acquisition (total pancreatectomy with auto islet transplantation (TPIAT) and partial pancreatectomy (PP), organoid generation, and pathologist fibrosis scoring (Fibrosis) information. For additional patient sample information see also Table S1, S2. Pancreas juice samples do not have histological sections for review and samples deemed histological failures could not be reviewed (N/A). (C) Mutational profile of patient samples as annotated through patient medical history for which PDOs were generated. Mutations in cystic fibrosis transmembrane conductance regulator (CFTR), serine protease inhibitor kazal type 1 (SPINK1), and protease serine 1 (PRSS1) are present in our patient samples representing some of the most common alterations in CP. Mutation - a mutation was annotated in patient medical history but the specific alteration was not reported. (D) Generation efficiency of patient derived organoids (PDOs); cumulative generation efficiency of 70.5%. (E) Representative images of human organoid cultures established from normal (hNP) or hCP after at least 4 passages. Scale bar 500 µm. (F) Whole mount immunofluorescence of PDOs stained with Hoechst (nuclei), EdU (pulsed for 25 hours), and CK19 (ductal marker). Scale bar 100 µm. (G)Quantification of EdU positive nuclei in a panel of NP and CP PDOs across two or three independent experiments and multiple passages. For representative images of immunofluorescence see Figure S1D.

**Table 1.**
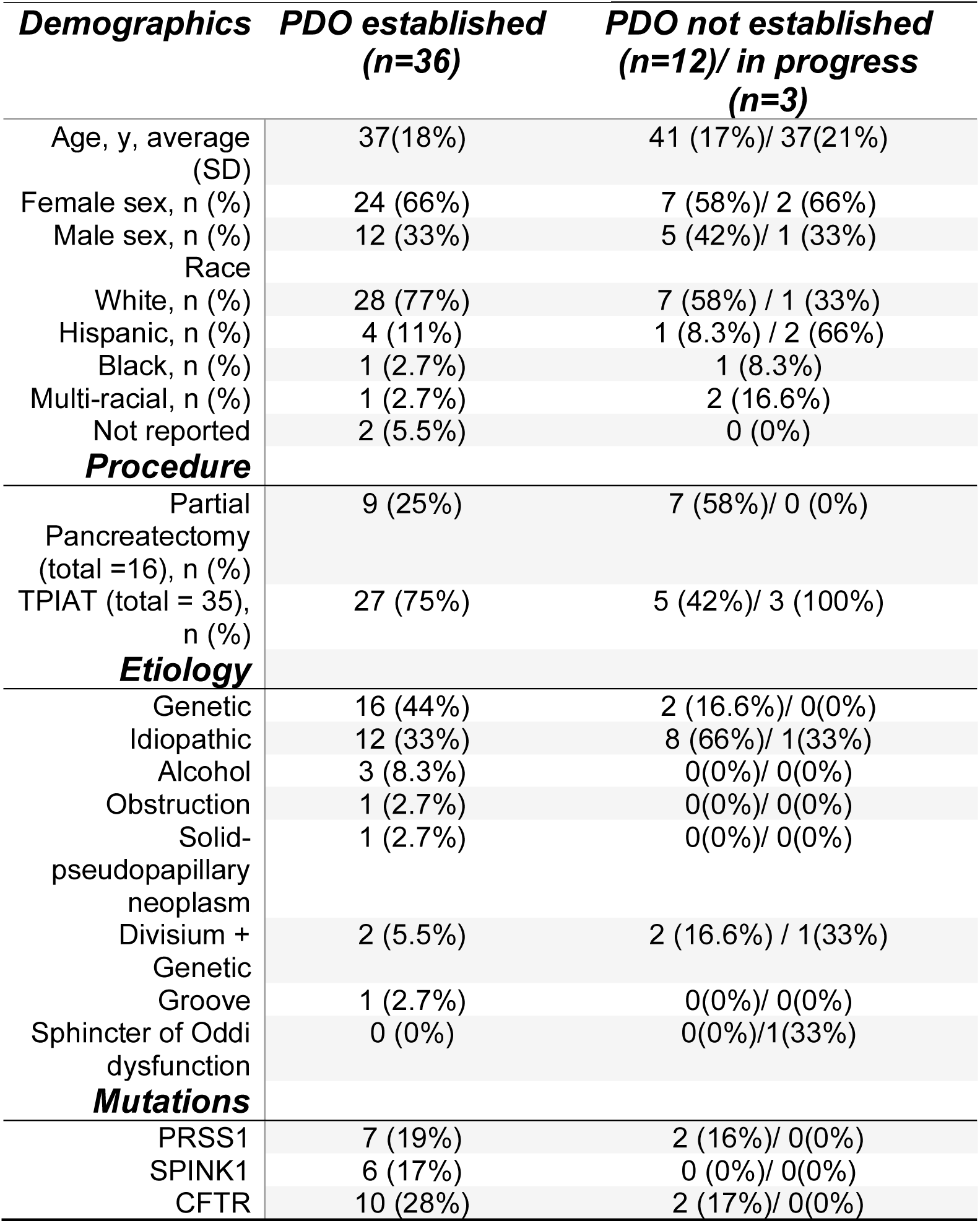
Characteristics of patients (n=51).

| **Demographics** | **PDO established**<br>**(n=36)** | **PDO not established**<br>**(n=12)/ in progress**<br>**(n=3)** |
| --- | --- | --- |
| Age, y, average<br>(SD) | 37(18%) | 41 (17%)/ 37(21%) |
| Female sex, n (%) | 24 (66%) | 7 (58%)/ 2 (66%) |
| Male sex, n (%) | 12 (33%) | 5 (42%)/ 1 (33%) |
| Race |  |  |
| White, n (%) | 28 (77%) | 7 (58%) / 1 (33%) |
| Hispanic, n (%) | 4 (11%) | 1 (8.3%) / 2 (66%) |
| Black, n (%) | 1 (2.7%) | 1 (8.3%) |
| Multi-racial, n (%) | 1 (2.7%) | 2 (16.6%) |
| Not reported | 2 (5.5%) | 0 (0%) |
| **Procedure** |  |  |
| Partial<br>Pancreatectomy<br>(total =16), n (%) | 9 (25%) | 7 (58%)/ 0 (0%) |
| TPIAT (total = 35),<br>n (%) | 27 (75%) | 5 (42%)/ 3 (100%) |
| **Etiology** |  |  |
| Genetic | 16 (44%) | 2 (16.6%)/ 0(0%) |
| Idiopathic | 12 (33%) | 8 (66%)/ 1(33%) |
| Alcohol | 3 (8.3%) | 0(0%)/ 0(0%) |
| Obstruction | 1 (2.7%) | 0(0%)/ 0(0%) |
| Solid-<br>pseudopapillary<br>neoplasm | 1 (2.7%) | 0(0%)/ 0(0%) |
| Divisium +<br>Genetic | 2 (5.5%) | 2 (16.6%) / 1(33%) |
| Groove | 1 (2.7%) | 0(0%)/ 0(0%) |
| Sphincter of Oddi<br>dysfunction | 0 (0%) | 0(0%)/1(33%) |
| **Mutations** |  |  |
| PRSS1 | 7 (19%) | 2 (16%)/ 0(0%) |
| SPINK1 | 6 (17%) | 0 (0%)/ 0(0%) |
| CFTR | 10 (28%) | 2 (17%)/ 0(0%) |

Most CP specimens were obtained following total pancreatectomy islet auto transplantation (TPIAT, 75%), and the remainder were from partial pancreatectomies (25%) (**Figures 1B, S1A-C, Table S1**). Pancreatectomy specimens were obtained within 2 hours post-resection with intact parenchyma but were often heavily cauterized (**Figure S1B-hCP1, hCP3, S1C-hCP10)**. TPIAT specimens were partially dissociated and then received typically 24-48 hours post-surgery, often with disrupted parenchyma. When sufficient intact tissue was available, a portion of the specimens were fixed for histology (**Figures 1A, S1A**). Histologic assessment of the primary specimens by a pathologist confirmed classic CP features, including parenchymal atrophy, fibrosis, and ductal abnormalities (**Table 1, S1, Figures 1A, S2A-B**). Fibrosis scoring and quantification of collagen deposition revealed substantial desmoplasia, with some specimens containing little to no remaining acinar epithelial tissues (**Table 1, S1, Figures 1B, S2A-B**).

Organoids were observed after 1-7 days in culture and could be passaged every 4-7 days, 12-25 times, similar to the passaging potential of normal pancreas (NP) organoids (**Figure S1D-E**). PDO generation efficiency was determined after 5 passages to exclude lines that did not expand. Generation of 3 PDO lines are still in process and have not yet reached 5 passages. Out of the 48 specimens that have reached evaluation criteria, 36 yielded PDO lines. We successfully generated organoids from 70% of CP specimens (**Table 1**, **Figure 1D**). Failures occurred due to a lack of viable epithelial cells when tissue was frozen upon arrival, extensively cauterized, or composed primarily of fibrosis (**Figure S2B, Table S1**). PDO generation efficiency was higher for TPIAT specimens (77%) relative to partial pancreatectomy samples (56%) (**Figure 1D**).

CP organoids displayed a consistent, homogenous cystic morphology with a single cell layer (**Figure 1E**). Organoid morphology, passage time, and passaging potential did not differ across CP etiologies (**Figure S1D-E**). CP PDOs are comprised of rapidly dividing ductal epithelial cells as demonstrated by incorporation of EdU and immunofluorescence (IF) for CK19 (**Figure 1F-G, S1F**).

### PDOs have high genetic concordance and reveal important and previously unannotated alterations

We assessed the genetic fidelity of PDOs by performing genomic analyses on matched primary and PDO specimens. Genomic alterations were compared between PDOs and primary tissue by whole genome sequencing (WGS), whole exome sequencing (WES), or targeted sequencing (**Figures 2, S3A, Table S1**). More than 9 million single nucleotide polymorphisms (SNPs) and almost 2 million insertions and deletions (indels) were compared by WGS, revealing an average 98% match rate with an intra correlation coefficient (ICC) of 0.96 between 14 pairs of PDOs and primary tissue (**Figures 2, S3A**). More than 80,000 SNPs and 7,000 indels were compared by WES, revealing an average 91% match rate with ICC of 0.83 between 6 pairs of PDOs and primary tissue (**Figures 2**, denoted by ^#^ in **S3A**). These data demonstrate the high genetic concordance between patient-matched primary tissue and PDOs. NP PDOs did not contain any pancreatitis-associated pathogenic mutations nor PDA-associated oncogenic alterations.

**Figure 2.**
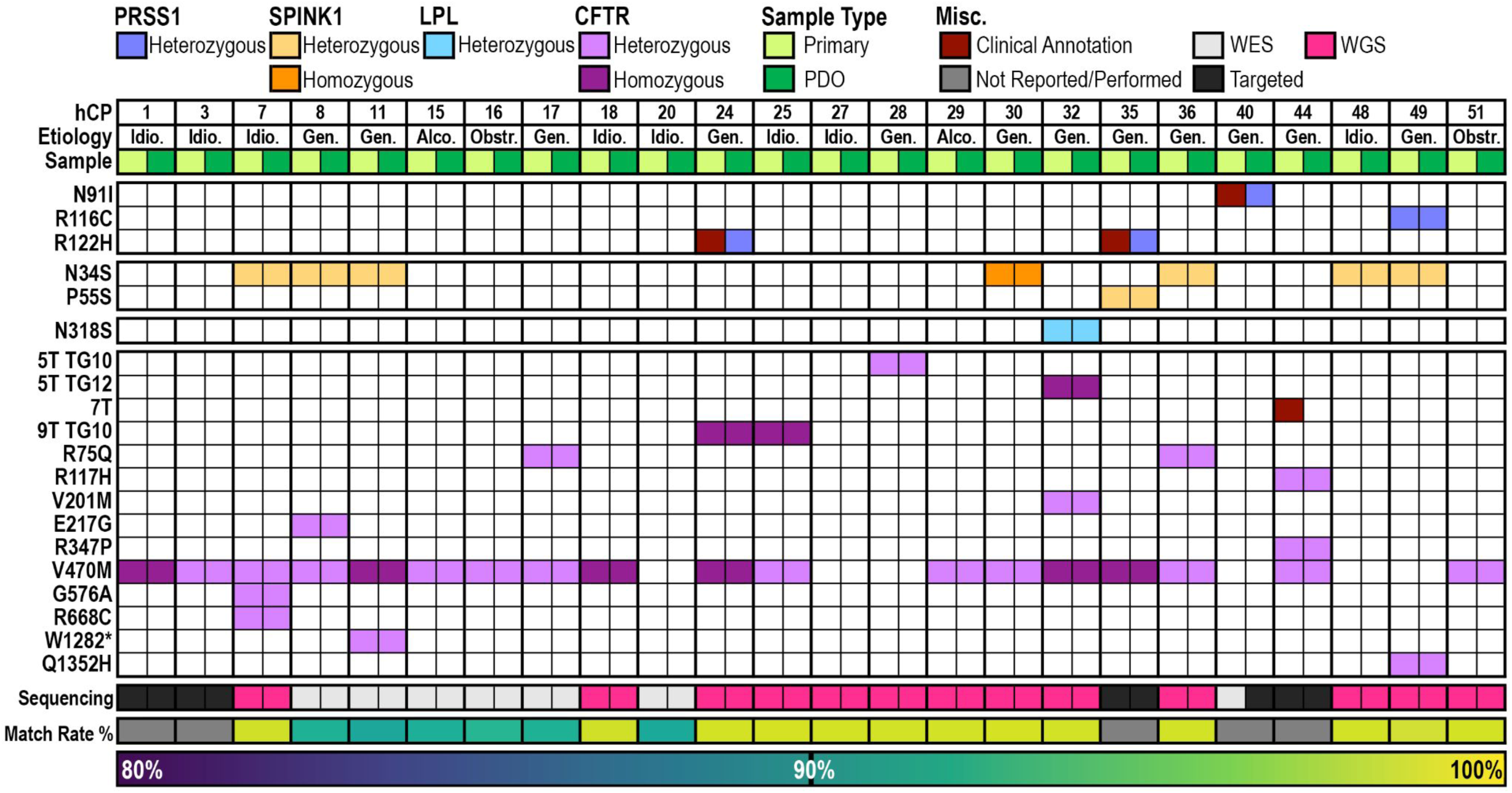
PDOs have high genetic concordance and reveal important and previously unannotated alterations. CP PDO and primary tissue coding sequence variants in PRSS1, SPINK1, LPL and CFTR determined from either WES (whole exome sequencing), targeted genetic sequencing, or WGS (whole genome sequencing). Genotype matches reference or is annotated as a heterozygous (Het) or homozygous (Homo) alteration as compared to reference. Clinical annotations from medical notes annotated if not found in PDO or primary tissue. Concordance is annotated for each pair of primary tissue and PDO as match rate (%). For concordance between unmatched pairs of primary tissue and PDO or PDOs see Figure S4A. For cystic fibrosis associated CFTR mutations annotated in our cohort see Figure S3B.

We then queried germline variants in 14 genes associated with hereditary pancreatitis. All clinically annotated PRSS1, SPINK1, and CFTR germline variants are retained in our PDOs (**Figure 2**). PRSS1 has five paralogs with 90-91% nucleotide similarity, resulting in reads aligning to multiple locations with low mapping quality scores (MapQ)^70,71^. The PRSS1 N91I mutation was called with high confidence (hCP49), while the R122H (hCP24, hCP35) and R116C mutations (hCP49) were identified with low MapQ scores (**Figure 2**). Mutations in SPINK1 were confirmed in both PDO and pancreatic tissue from hCP08, hCP11, hCP30, hCP36, and hCP49. Interestingly, in 2 PDOs from patients with idiopathic CP with no previous genetic testing, we found SPINK1 N34S mutation (hCP07, hCP48) (**Figure 2**). In addition, we identified an unannotated SPINK1 P55S mutation (hCP35) and LPL N318S mutation (hCP32), which is associated with hyperlipidemia^72^ and hereditary CP^73^ (**Figure 2**).

Intronic variants of CFTR are common^30^ and were confirmed in agreement with patient genetic testing results (hCP25, hCP28, and hCP32) (**Figure 2**). 9T variants are included in our assessment because their impact on exon 9 skipping in the pancreas in unknown despite minimal effect in nasal epithelium^74^. We confirmed the PDOs recapitulate coding sequence variants of CFTR, including R75Q, R117H, E217G, R347P, W1282*, and Q1352H mutations (**Figure 2**)^75–81^. Additionally, 75% of the CP PDO cohort harbors the CFTR V470M variant, which has conflicting reports of pathogenicity in association with CP and is likely a passenger mutation^75,80–84^. Additional G576A and R668C CFTR mutations were detected in hCP07, which have been found to be co-mutated in hereditary pancreatitis^85–87^. Also, a mild V201M CFTR variant was found in hCP32^88^. CFTR mutations associated with hereditary CP are distinct from cystic fibrosis and often do not overlap^25–32^. None of the cystic fibrosis associated CFTR mutations (CFTR2 database, n=1092) are found in the CP PDO cohort (**Figure S3B**).

Importantly, CP is a risk factor for developing pancreatic cancer and some hereditary CP patients have up to a 55% lifetime risk. The PDO platform enables identification of potential somatic oncogenic variants that are present in a small percentage of epithelial cells that would be obscured by the low epithelial cellularity of primary tissues. Here, we identified oncogenic KRAS G12D mutation in hCP30 with an allele frequency of 0.21 that was not detected in the primary specimen. We also found oncogenic TP53 R175H mutation in hCP28 at an allele frequency of 0.05 that was not detected in the primary specimen. These data suggest that the CP PDO models may be useful in identifying patients at higher risk of progressing to PDA.

### Gene expression analyses demonstrate distinct cell signaling programs in CP organoids

To understand the specific programs differentially regulated in CP PDOs, we performed RNA-seq on 24 CP, 9 NP, and 4 normal adjacent to tumor (NAT) PDOs. Ductal lineage identity is maintained by all PDOs with no evidence of non-epithelial cell contamination (**Figure S3C**), concordant with positivity for the ductal epithelial marker CK19 by IF (**Fig. 1F**). CP PDOs have a distinct transcriptomic profile as compared to NP and NAT PDOs (**Figures 3A-C**). Gene set enrichment analyses (GSEA) reveal that CP PDOs are enriched in gene sets associated with inflammatory response, TNFA signaling via NFKB, KRAS signaling, Epithelial to Mesenchymal Transition (EMT), and TGF beta signaling while having depressed Oxidative Phosphorylation relative to NP and NAT PDOs (**Figure 3D**). Compared to NP and NAT PDOs, CP expressed higher levels of fibrosis and inflammatory cytokine associated genes (e.g., *COL1A1*, *TGFB2*, *IL1A*, *CSF1*) (**Figure 3E**), suggesting that CP PDOs maintain the fibrosis and inflammatory programs present in CP patients^89^.

**Figure 3.**
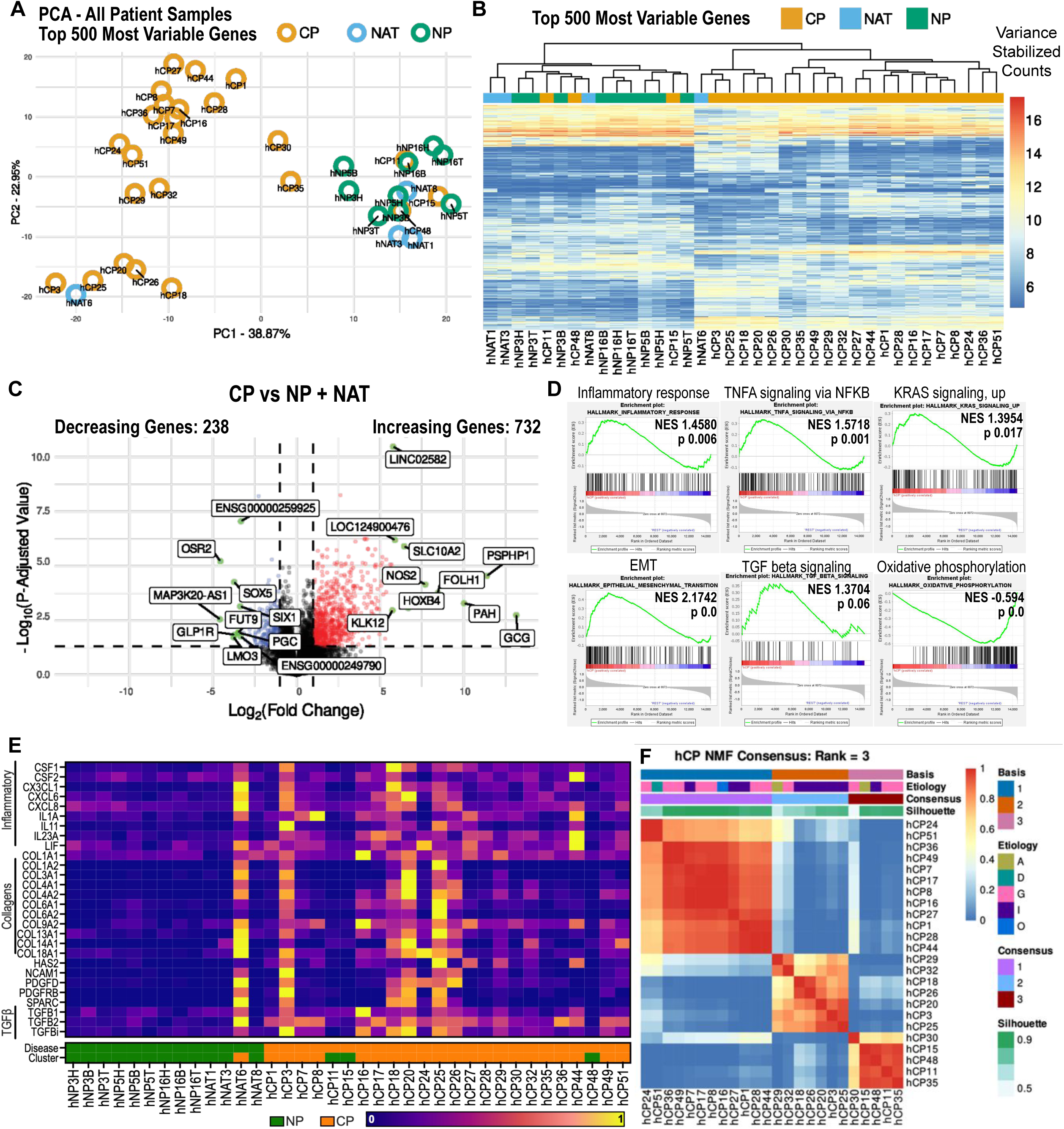
Gene expression analyses demonstrate distinct cell signaling programs in CP organoids. (A) Principal component analysis (PCA) of top 500 most variable genes from organoids isolated from NP, normal adjacent to tumor (NAT), and CP samples. (B) Top 500 variable genes from variance stabilized gene expression in hCP, hNAT, and hNP samples. Hierarchical clustering separates NP and NAT from most CP organoids. (C) Volcano plot of CP versus NP and NAT samples. (p value <0.05, |log2fold change| > 1 colored). Top 10 enriched or depleted genes annotated. (D) Gene set enrichment analysis of CP PDOs versus NP and NAT PDOs. (E) Relative normalized transcript expression of select inflammatory, collagens, and TGFb genes from RNAseq. Sample disease origin and cluster are denoted. For more information on cell lineage genes see Figure S4. (F) Clustering using non-matrix factorization (NMF) defines three distinct subtypes of hCP PDO cultures; basis 1, 2, and 3. Patient etiology (A-alcoholic, D-divisum + groove, G-genetic, I-idiopathic, O-obstruction), silhouette, and consensus is indicated. For NMF rank survey see Figure S4.

Principal component analysis of CP PDOs suggested the presence of 3 distinct clusters (**Figure S4A**). The PDO transcriptomes were independently clustered using nonnegative matrix factorization (NMF)^63,64^, revealing 3 stable subtypes in CP PDO cultures (basis 1-3) (**Figures 3F**, **S4B**). The molecular subtypes identified using this approach do not cluster according to etiology (**Figure 3F**), suggesting distinct subtypes independent of TIGAR-O classification. We performed GSEA on each CP subtype relative to NP and NAT PDOs (**Figure S4C**). The basis 1 subtype is depleted in oxidative phosphorylation and enriched in EMT and apoptosis gene sets. The apoptosis gene set includes proteins that involve cellular stress response pathways, such as glutathione peroxidase 3 (GPX3). The basis 2 subtype is enriched in E2F targets, MYC targets, and myogenesis. The basis 3 subtype is enriched in E2F targets. All subtypes trend towards enrichment of the inflammatory response gene set.

To validate these subtypes in matched primary tissue specimens, we identified the genes that were differentially expressed between the 3 subtypes of CP and have the highest Kim-score, and therefore drive these distinct clusters (**Figure S4D**). Subtype defining genes were evaluated by RNA *in situ* hybridization (**Figure S5A**). We found that a patient specimen from the basis 1 subtype PDO hCP1 expressed the highest levels of GPX3 as expected. The patient matched specimen for the basis 2 subtype PDO hCP3 exhibited the highest level of aquaporin 1 (AQP1). The basis 3 subtype was defined by BPI-family member B 1 (BPIFB1), which exhibited the highest levels in patient specimen from the basis 3 subtype PDO hCP48. This indicates that the genes driving CP PDO transcriptional subtypes can also be detected in the corresponding primary tissue.

Transcriptomic analyses on primary CP tissues are challenging due to the high degree of fibrosis and the high levels of RNases. We further explored the expression of subtype-defining genes using single-nucleus RNA sequencing (snRNA-seq) data from human NAT and CP samples^69^. In the ductal cell cluster, we observed that subtype defining genes were expressed in primary pancreatic ductal epithelium (**Figure S5B-C**), but insufficient sample and cell numbers made identification of subtypes in these data challenging. These findings confirm that the genes driving transcriptional subtypes in the CP PDOs are present in an independent patient cohort.

Additional analysis comparing the transcriptomes of our CP PDO cohort to human pancreatic cancer organoids revealed genes that are uniquely expressed in either disease (**Figure S5D**). This along with the discovery of somatic oncogenic variants in 2 of our CP PDOs demonstrate that this model can be used to study patients with heightened risk of cancer and to identify early detection biomarkers.

### CP organoids recapitulate protein expression from primary tissue specimens and exhibit elevated inflammatory and mitogenic signaling

Immunohistochemistry (IHC) and immunoblotting (IB) were used to evaluate proteins and glycan levels associated with pancreatitis, as well as mitogenic and inflammatory signaling. CP and NP PDOs exhibited protein alterations consistent with matched patient specimens (**Figures 4A-B**). We previously found that up to ∼90% of adult CP has local elevation of CA19-9^90^. Patient specimens had a range of CA19-9 that matched CP PDOs (**Figures 4A-B**). Activation of EGFR has been reported in human CP^91^. A subset of CP PDOs have elevated levels of phospho-EGFR and downstream effectors AKT and ERK relative to NP PDOs (**Figure 4A**), suggesting there are “mitogenic” and “non-mitogenic” CP subtypes. NP PDOs primarily express mature CFTR by IB (upper band, C-band, 150-160KD^92^) with very little immature CFTR (lower band, B-band, 135-140KD^92^) (**Figures 4A, S6A**). Of those with CFTR mutations, hCP11 and hCP44 had low levels of mature CFTR by IB, which was replicated in the primary patient specimen IHC (**Figures 4A-B**). This corresponds with what is known about the pre-mature stop codon mutation present in hCP11 and compound CFTR R347P; R117H mutation in hCP44. These mutations have been reported to lead to major folding defects, and thus, as expected, do not produce mature CFTR^79^. E217G and R75Q CFTR mutations found in hCP08 and hCP17, respectively, impact anion channel function^75,92^, but not protein maturation. These two lines have mature CFTR, as predicted. hCP15, a non-hereditary case attributed to alcohol and hypothyroidism, also demonstrated loss of mature CFTR protein despite having a benign V470M mutation (data not shown). Decreased CFTR expression and function has been shown to occur secondary to alcohol^93,94^, tobacco^95^, and hypothyroidism^96^. These findings suggest that CFTR modulators that increase protein activity may help even CP patients without pathogenic CFTR mutations. Carbonic anhydrase 2 (CA2), which catalyzes the conversion of CO_2_ into HCO_3_^-^, was decreased in most CP PDOs when compared to NP, pointing to another possible mechanism through which HCO_3_^-^ secretion may be impaired in CP (**Figure 4A**).

**Figure 4.**
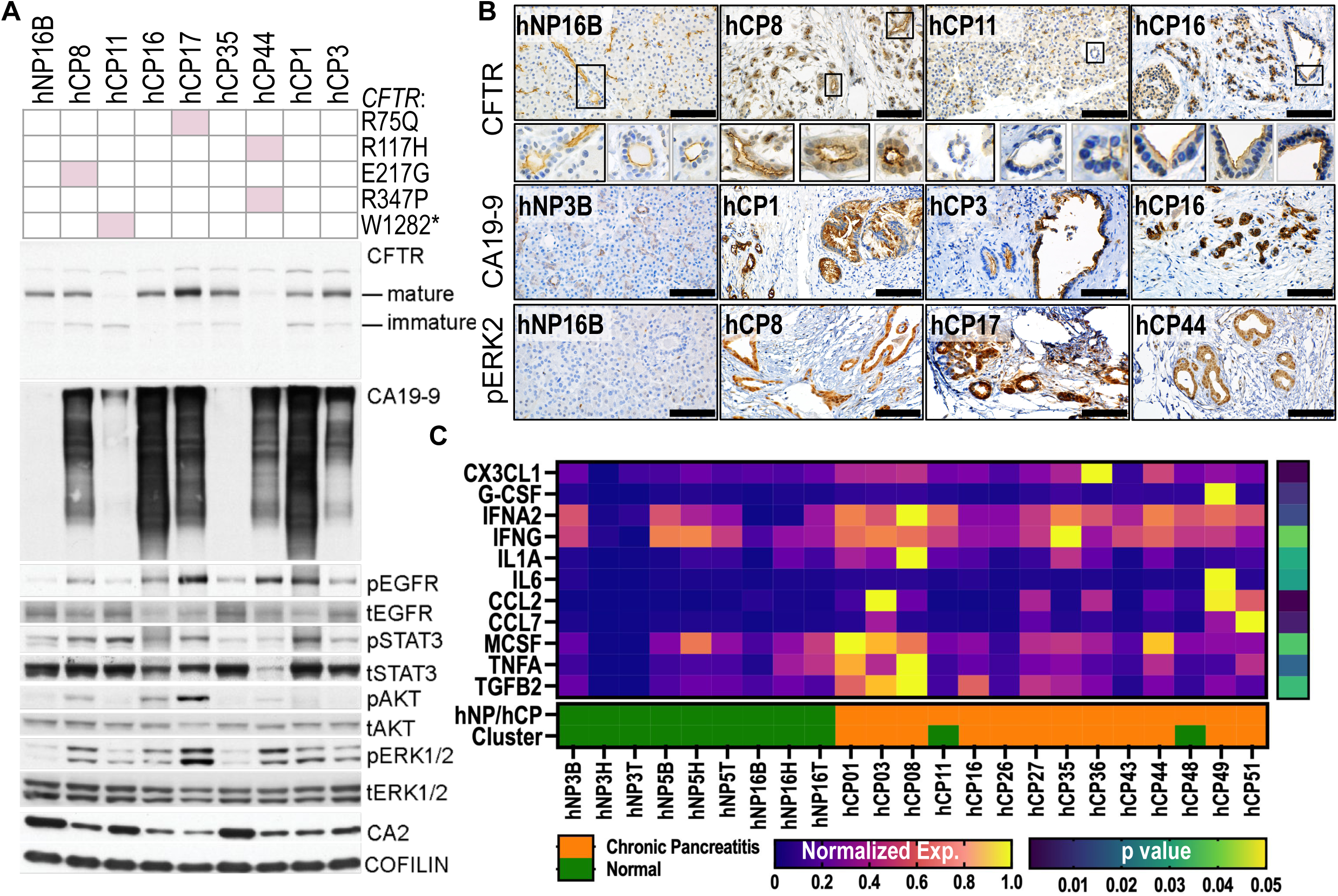
CP organoids recapitulate protein expression from primary tissue specimens and exhibit elevated inflammatory and mitogenic signaling. (A) Immunoblot of PDOs generated from NP and CP samples. CFTR mutations annotated for CP samples. For normalized mature CFTR expression across cohort see Figure S7A. (B) Immunohistochemistry (IHC) staining for cancer antigen 19-9 (CA19-9), signal transducer and activator of transcription 3 phosphorylation (pSTAT3), mitogen-activated protein kinase 1 phosphorylation (pERK2), and cystic fibrosis transmembrane conductance regulator (CFTR) of NP and CP samples. Scale bar 100 µm. (C) Normalized secreted protein expression of inflammatory cytokines in NP and CP PDOs. Average concentration of two technical replicates per sample. P-value determined by Mann-Whitney test. Sample type and cluster is shown. For additional cytokines assayed see Figure S7B.

To identify fibroinflammatory mediators in CP PDOs at the protein level, we evaluated conditioned medium from 14 CP and 9 NP PDOs. IL1A, TGFB2, CX3CL1, and other fibroinflammatory secreted factors were elevated in CP PDOs and reflected the changes observed in gene expression and published changes in CP patients^97,98^ (**Figure 4C, S6B**), demonstrating the ability of CP PDOs to model inflammatory changes seen in patients. Together, these data support that PDOs retain key molecular features of CP, including alterations in cell signaling proteins.

### CFTR function in CP organoids is restored upon pharmacologic modulation

The forskolin-induced swelling (FIS) assay was developed to measure CFTR dysfunction and previously deployed in intestinal and rectal organoids from cystic fibrosis patients to predict response to CFTR modulators^39,40^. Intestinal current measurements and FIS activity were tightly correlated, demonstrating that FIS assays can serve as a surrogate for more complex electrophysiological measurements^40^. We optimized the FIS assay in murine and human NP PDOs (**Figure 5A**). Vehicle treated organoids had no response, but forskolin or CFTR potentiator treatment caused organoid swelling (**Figures 5B-F, S7A-C**). Swelling was inhibited when mouse and human NP organoids were treated with CFTR inhibitors (CFTRi-172, PPQ-102) or a PKA inhibitor (H-89), demonstrating that swelling is dependent on forskolin induced activation of CFTR via PKA (**Figures 5D-F, S7B**). Mouse normal organoids served as a positive control while testing the CFTR function of CP PDOs (**Figure S7A-B)**. Half of the CP PDOs evaluated (n=4 out of 8) were FIS impaired across multiple experimental replicates (**Figure 5G**, **S7D**).

**Figure 5.**
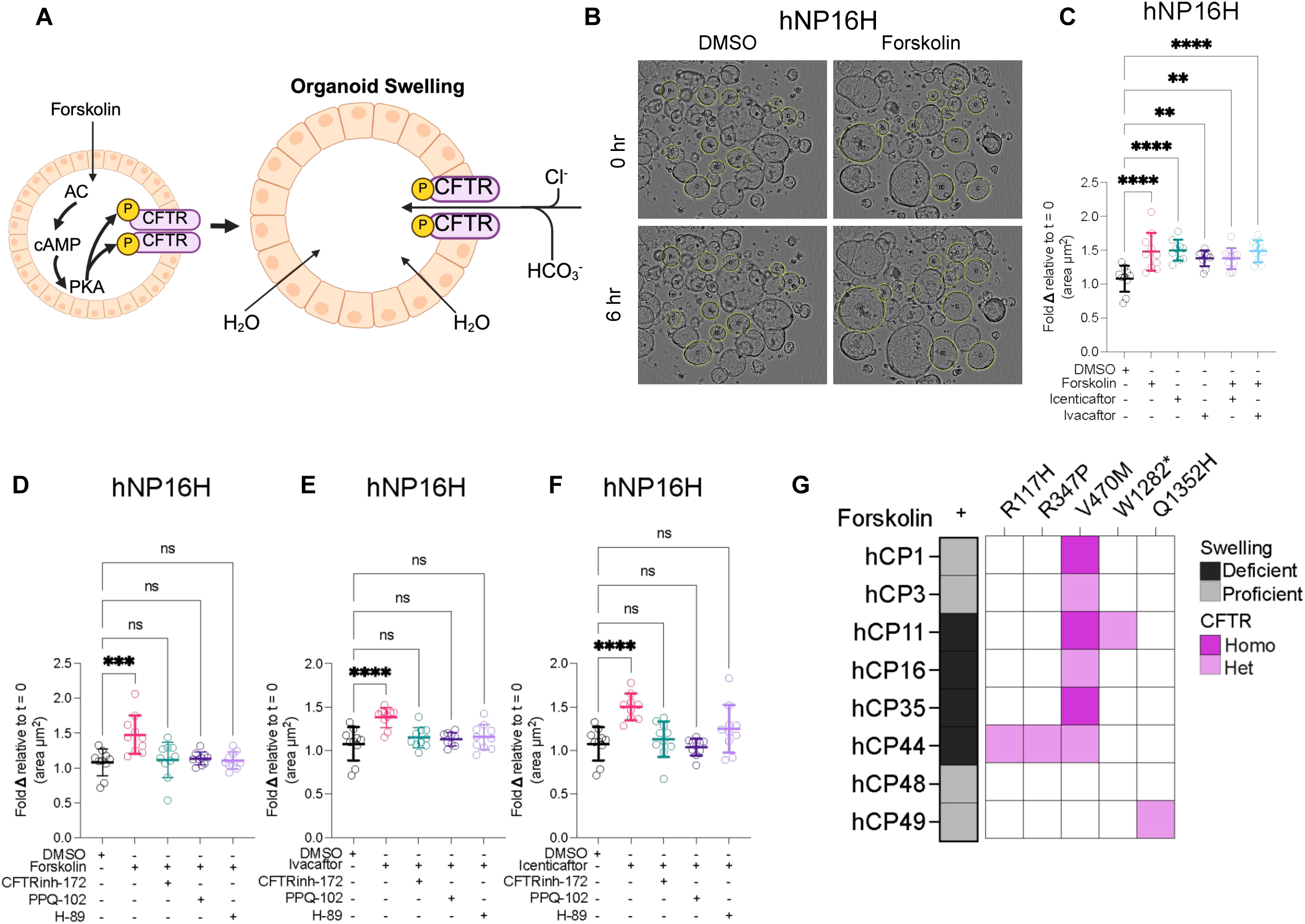
Forskolin induced swelling of CP PDOs is CFTR dependent. (A) Schematic of forskolin induced swelling assay. Created in BioRender. Osorio Vasquez, V. (2025) https://BioRender.com/oahkr5p (B)Representative brightfield images of organoids used for the quantification of forskolin induced swelling assay. Upto 10 organoids are selected per condition. (C) Forskolin induced swelling (FIS) assay of hNP organoids with forskolin, CFTR potentiators (icenticaftor or ivacaftor), or a combination of both for 6 hours. (D) Forskolin induced swelling (FIS) assay of hNP organoids treated with forskolin for 6 hours and pre-treated with CFTR inhibitors (CFTRi-172, PPQ-102), or protein kinase A inhibitor (H-89) for 2 hours. Swelling is inhibited with the addition of CFTR and PKA inhibitors. (E) Forskolin induced swelling (FIS) assay of hNP organoids treated with ivacaftor for 6 hours and pre-treated with CFTR inhibitors (CFTRi-172, PPQ-102), or protein kinase A inhibitor (H-89) for 2 hours. Swelling is inhibited with the addition of CFTR and PKA inhibitors. (F) Forskolin induced swelling (FIS) assay of hNP organoids treated with icenticaftor for 6 hours and pre-treated with CFTR inhibitors (CFTRi-172, PPQ-102), or protein kinase A inhibitor (H-89) for 2 hours. Swelling is inhibited with the addition of CFTR and PKA inhibitors. (G) FIS assay of hCP organoids treated with forskolin for 6 hours. Data represented as swelling deficient or proficient compared to DMSO control at 6 hours. CFTR mutations are shown for each sample. Data represented as mean ± SD for C, D, E. Concentrations for forskolin treatments 20 µM, CFTR potentiators (ivacaftor, icenticaftor) or correctors (elexacaftor, lumacaftor) 10 µM for FIS assays. Data represented as swelling deficient or proficient compared to DMSO control of at least 2 independent experiments. ETI treatment concentrations are 3 µM for each compound.

Most CFTR wild type PDOs (hCP1, hCP3, hCP48, hCP49) swelled in response to forskolin treatment as expected (**Figure 5G**). Unexpectedly, hCP16, which is CFTR wildtype and exhibits normal CFTR protein levels (**Figure 4A**), is FIS deficient (**Figure 5G, S7D**). We hypothesized that organoids deficient in FIS could be rescued by treatment with CFTR potentiators, which increase the basal activity of CFTR (**Figure 6A**). Treatment of hCP16 with CFTR potentiators did not restore swelling (**Figure S7D**), suggesting that the dysfunction in FIS in this PDO is independent of CFTR.

**Figure 6.**
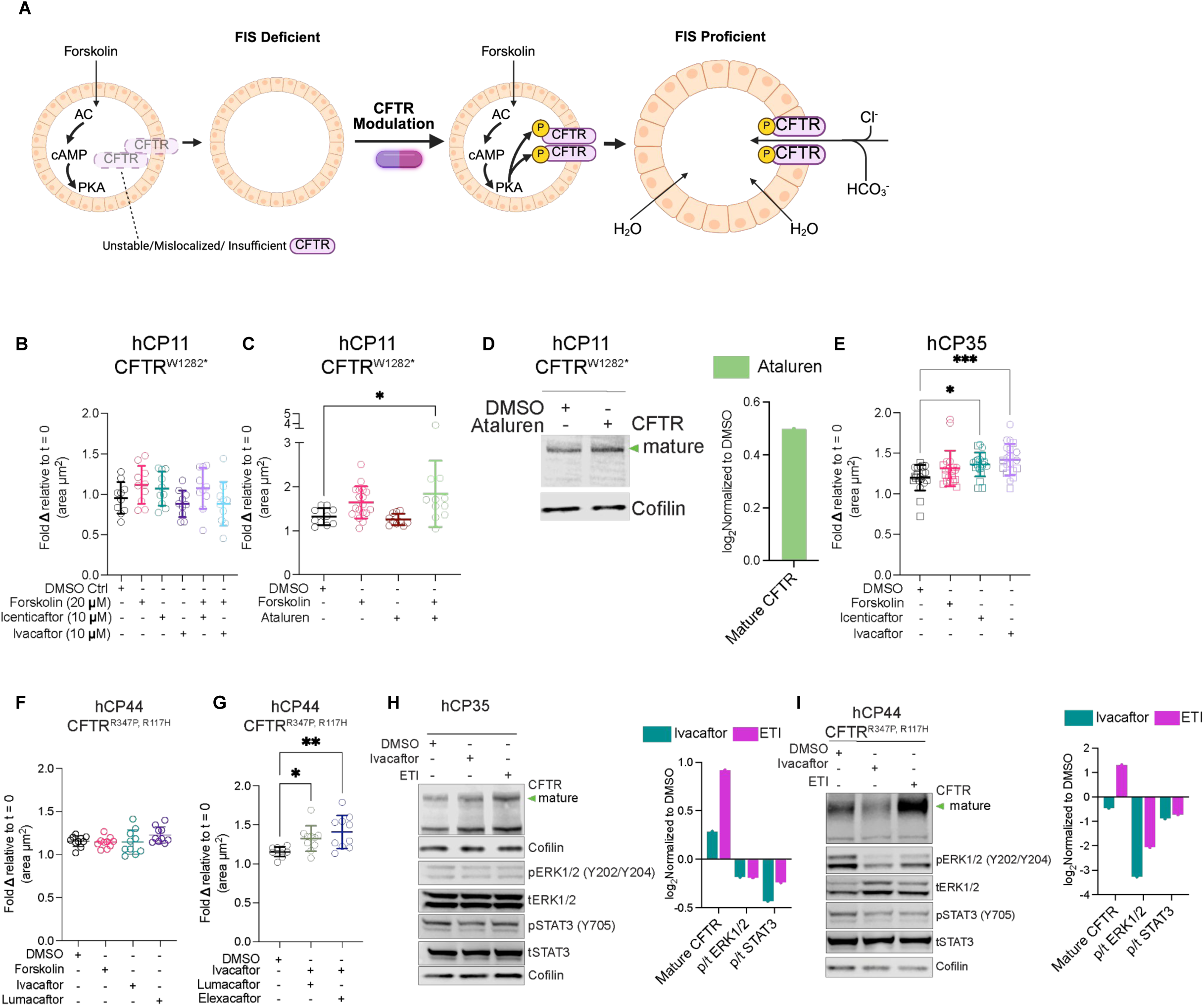
CFTR function in CP organoids is restored upon pharmacologic modulation. (A) Schematic of rescuing impaired forskolin induced swelling in PDOs. Created in BioRender. Osorio Vasquez, V. (2025) https://BioRender.com/a9tuea3 (B) hCP11 (CFTR W1282*) is FIS deficient upon treatment with forskolin or CFTR potentiators (icenticaftor or ivacaftor) for 6 hours. (C)Treatment with CFTR readthrough compound (Ataluren-10 µM) and 6 hour treatment with forskolin in hCP11 (CFTR W1282*) induces swelling. (D) IB of mature CFTR expression upon 48 hour Ataluren (10 µM) treatment in hCP11 (CFTR W1282*). Quantification of IB directly adjacent. (E) FIS is rescued upon 6 hour CFTR potentiator (Ivacaftor and Incenticaftor) and forskolin treatment in hCP35 PDOs. Data representative of at least 2 independent experiments. (F) hCP44 (CFTR R347,R117H) organoids did not swell after 6 hours of treatment with DMSO, forskolin, CFTR potentiator (Ivacaftor), or CFTR corrector (lumacaftor). (G) hCP44 (CFTR R347,R117H) organoids swelled in response to combination CFTR potentiator (Ivacaftor) and CFTR corrector (lumacaftor or elexacaftor) treatment after 6 hours. (H) Mature CFTR expression is increased in hCP35 organoids treated with CFTR potentiator (Ivacaftor) or further increased with ETI (CFTR correctors-Elexacaftor and Tezacaftor, CFTR potentiator-Ivacaftor) for 48 hours. pERK2 and pSTAT3 expression decreases upon ETI treatment. Cofilin is used as loading control. Representative blot of 3. Quantification of IB relative to DMSO adjacent. (I) Mature CFTR expression increases upon ETI treatment for 48 hours (CFTR correctors-Elexacaftor and Tezacaftor, CFTR potentiator-Ivacaftor) in hCP44 (CFTR R347,R117H) organoids. pERK1/2 and pSTAT3 is signaling is decreased upon ETI treatment. Cofilin is used as loading control. Quantification of IB relative to DMSO adjacent. Data represented as mean ± SD for C, D, E. Concentrations for forskolin treatments 20 µM, CFTR potentiators (ivacaftor, icenticaftor) or correctors (elexacaftor, lumacaftor) 10 µM for FIS assays. Data represented as swelling deficient or proficient compared to DMSO control of at least 2 independent experiments. ETI treatment concentrations are 3 µM for each compound.

hCP11 did not swell, which can be attributed to the lack of mature CFTR due to a premature stop codon mutation (**Figures 4A**)^99^. Given the lack of mature CFTR protein, treatment with CFTR potentiators did not restore swelling (**Figure 6B**). However, treatment with the read through agent, Ataluren, was able to restore FIS response (**Figure 6C**) and increase levels of mature CFTR (**Figure 6D**). These data demonstrate that we can successfully achieve genotype-driven precision therapeutics in CP PDOs.

hCP35 is CFTR wildtype and exhibits normal CFTR protein levels. Unexpectedly, hCP35 had a limited FIS response (**Figures 5G, 6E**). Swelling was restored by treatment with CFTR potentiators, Ivacaftor and Icenticaftor (**Figure 6E**). This demonstrates ability of potentiators to increase CFTR function in PDOs even with wild type CFTR.

hCP44 was also deficient in FIS (**Figure 6F, 6I**), which was expected given the CFTR R347P, R117H mutations that impair protein folding and anion transport^100^ and because the low levels of mature CFTR protein (**Figure 4A**). The protein folding defect can be ameliorated with CFTR correctors. This class of drugs increases the correct folding and plasma membrane localization of CFTR and the combination of correctors and potentiators is FDA approved for the treatment of cystic fibrosis (e.g., Orkambi: Ivacaftor, Lumacaftor)^101^. hCP44 FIS was rescued by treatment with a combination of correctors and potentiators (Orakambi, and Ivacaftor + Elexacaftor) (**Figure 6G**), but not monotherapies (**Figure 6F**). This demonstrates that CP PDO models can used to identify the most effective combinatorial treatment strategies in the background of complex allelic combinations.

We tested if treatment with CFTR correctors and potentiators increases mature CFTR protein levels in FIS deficient hCP35 and hCP44. Single agent treatment of hCP35 with the potentiator Ivacaftor marginally increased mature CFTR, which was further increased by ETI treatment (**Figure 6H**). As expected from our FIS results, ETI treatment increases mature CFTR protein levels in hCP44 (**Figure 6I**). Importantly, treatment of hCP44 with single agent CFTR potentiator caused a decrease in mature CFTR protein levels (**Figure 6I**). This is in line with previous findings that Ivacaftor treatment can interfere with CFTR stability in the background of certain mutations, such as the R347P, R117H alterations found in hCP44^102^. These findings stress the importance of investigating CFTR modulator response in patient-specific contexts, especially in the setting of complex allelic combinations. Altogether these data indicate that patients with and without pathogenic *CFTR* alterations could benefit from ETI treatment to increase mature CFTR expression. Treatment with ETI in both hCP35 and hCP44 also led to a decrease in pERK and pSTAT3 levels (**Figures 6H-I**), suggesting that restoration of CFTR function can decrease mitogenic and inflammatory signaling. Our findings suggest that restoring normal ductal function can contribute to the resolution of inflammation.

## Discussion

Surgical samples from chronic pancreatitis patients are rare, making this CP PDO biobank a unique resource. Importantly, while our PDO biobank encompasses the major etiologies of CP, the patients are a highly selected cohort that required surgical management. In the future, this limitation may be addressed through generation of CP PDOs from pancreatic juice, which we accomplished in 2 cases and is a more accessible sample source.

The identification of pathogenic germline alterations in 3 patients diagnosed with idiopathic CP suggests that genetic testing may have broader benefit in CP clinical care. Two of the CP PDOs also contain oncogenic mutations in TP53 or KRAS, known drivers of pancreatic ductal adenocarcinoma. This is expected given the high rate of sporadic KRAS mutations that have been found in autopsy studies of the normal human pancreas^103^. Work has also shown that even in the context of wild type KRAS, TP53 mutation promotes injury in a non-cancer context^104^. CP PDO models will facilitate our understanding of which patients are at risk of severe disease and progression to PDA. Identification of the KRAS and TP53 mutations were uniquely possible from analysis of the PDOs and not the tissue. Further work is needed to explore how CP PDOs can be leveraged to model the transition from a chronic inflammatory state to invasive carcinoma.

CP is primarily classified by etiology, but our transcriptomic analyses reveal molecular subtypes that do not correlate with etiology, suggesting that distinct cell signaling changes drive CP subtypes. The validation of these subtypes remains challenging due to the limited availability of these surgical tissue specimens and the high abundance of RNases and proteases in the pancreas. Future work will focus on creating primary CP specimen data resources for corroboration of these findings and for investigation of clinical correlates. These subtypes offer an initial framework to explore potential signaling pathways driving CP pathogenesis.

Surprisingly, the CP PDOs maintain a fibroinflammatory gene and protein expression signature despite being removed from the microenvironment. These data suggest that ductal cells can contribute to maintenance of the fibroinflammatory program and opens several lines of questioning. When are these ductal gene expression programs established and how are they maintained? Is the intrinsic epithelial fibroinflammatory program sufficient to cause CP and promote PDA initiation? Would reversing this program restore pancreatic function? CP PDOs enable exploration of these important questions at unprecedented depth as well as the appropriate physiologic context.

The use of PDOs for functional precision medicine has been previously established in pancreas cancer^43,46,50,52^. Here, we show that CP PDOs can be similarly used as a platform for studying CFTR function in different complex genetic contexts and delineating treatment response. Our study shows that CP PDOs are commonly deficient in FIS. This deficiency can be effectively reversed by treatment with FDA-approved CFTR modulators, suggesting potential targeted therapeutic strategies for the treatment of CFTR-mutant CP. An important consideration when testing CFTR modulators to treat CP includes organ context and species. CFTR mutations in cystic fibrosis patients primarily impacts chloride secretion whereas in the pancreas, it contributes substantially to bicarbonate transport. This highlights the importance of investigating CP CFTR mutations in pancreatic cell types. In addition, mutant mouse CFTR does not respond to human CFTR modulators^41^. This is further complicated by reduced levels of bicarbonate in mouse pancreatic juice (50 mM) relative to human (140 mM)^37^, suggesting a lower level of reliance on bicarbonate in the mouse pancreas. Finally, as evidenced in our cohort, the genetics of CP is complex. There are more than 2,000 different CFTR mutations and limited information regarding drug response to most of these alterations, especially those associated with hereditary CP. Even in the setting of the same pathogenic alterations, patient response to CFTR modulators is very heterogeneous, highlighting the need for precision medicine approaches^35–40^. The physiologically relevant context of human CP PDOs enables direct investigation of genotypic relationships to therapeutic responses. We further demonstrate CFTR dysfunction in CP PDOs with wildtype CFTR that can be restored with potentiator treatment, raising the possibility that this treatment approach could be widely applicable to a large patient population. Our work also highlights the utility of restoring normal homeostatic function to reduce inflammatory signaling as evidenced by decreased STAT3 activation following CFTR modulator treatment. Altogether, we demonstrate the power of our PDO platform to study the underlying molecular pathogenesis of human CP and pave the way for precision therapeutics.

## Acknowledgements

The authors would like to extend our deepest gratitude to all the patients who generously consented to participate in this study. The authors would like to thank all the members of the Engle laboratory, the UCSD Biorepository and Tissue Technology Shared Resource (BTTSR), the UCSD Department of Surgery, the University of Minnesota Departments of Surgery and Pediatrics, April Williams, Ling Ouyang, and Tzu-when Wang at the NGS Core Facility of the Salk Institute, Elsa Molina at the Single-Cell and Spatial Omics core of the Salk Institute, Daniela Boassa, Sammy Weiser Novak and Elsie Quansah at the Biophotonics Core Facility of the Salk Institute, and the Lifesharing Donate Life organization.

## Funding Statement

V.O.V is supported through the Salk Institute Cancer Training Grant (T32CA009370). J.C.L. was supported through the Salk Institute Cancer Training Grant (T32CA0093790). This work was supported by the Razavi Newman Integrative Genomics and Bioinformatics Core Facility of the Salk Institute with funding from NIH-NCI CCSG: P30 CA01495, NIH-NlA San Diego Nathan Shock Center P30 AG068635, and the Helmsley Trust. This work was supported by the Waitt Advanced Biophotonics Core Facility of the Salk Institute with funding from NIH-NCI CCSG: P30 CA01495, NIH-NlA San Diego Nathan Shock Center P30 AG068635, and the Waitt Foundation. This work was supported by the NGS Core Facility of the Salk Institute with funding from NIH-NCI CCSG: P30 CA01495, NIH-NlA San Diego Nathan Shock Center P30 AG068635, the Chapman Foundation and the Helmsley Charitable Trust. The Biorepository and Tissue Technology Shared Resource is supported by a National Cancer Institute Cancer Center Support Grant (CCSG P30CA23100) and by recharges from users. This research program was supported by Mission Cure Capital, LLC, the Lustgarten Foundation, the Conrad Prebys Foundation, The Paul M. Angell Family Foundation, Curebound (20DG11), and by the Helen McLoraine Developmental Chair.

## Land Acknowledgement

The Salk community holds great respect for the land and the original people of the area where our campus is located. The Institute is built on the territory of the Kumeyaay Nation. Today, the Kumeyaay people continue to maintain their political sovereignty and cultural traditions as vital members of the San Diego community. We acknowledge their tremendous contributions to our region and thank them for their stewardship.

## Declaration of Interests

J. Gibson, and D. Whitcomb are founders of Ariel Precision Medicine. D. Whitcomb is a member of its scientific advisory board at Ariel Precision Medicine. P. Greer is an employee of Ariel Precision Medicine.

---

**Table S1.**
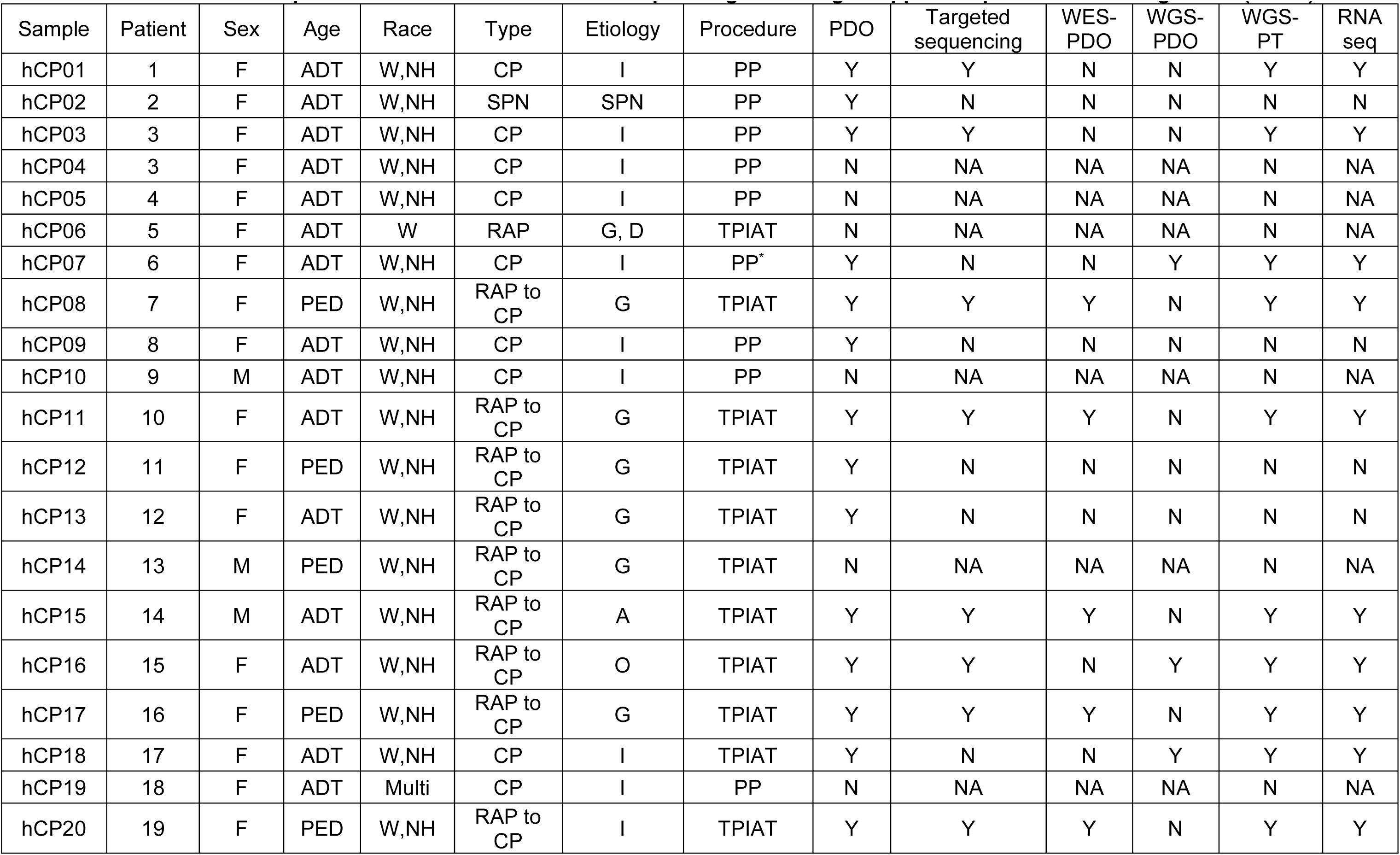
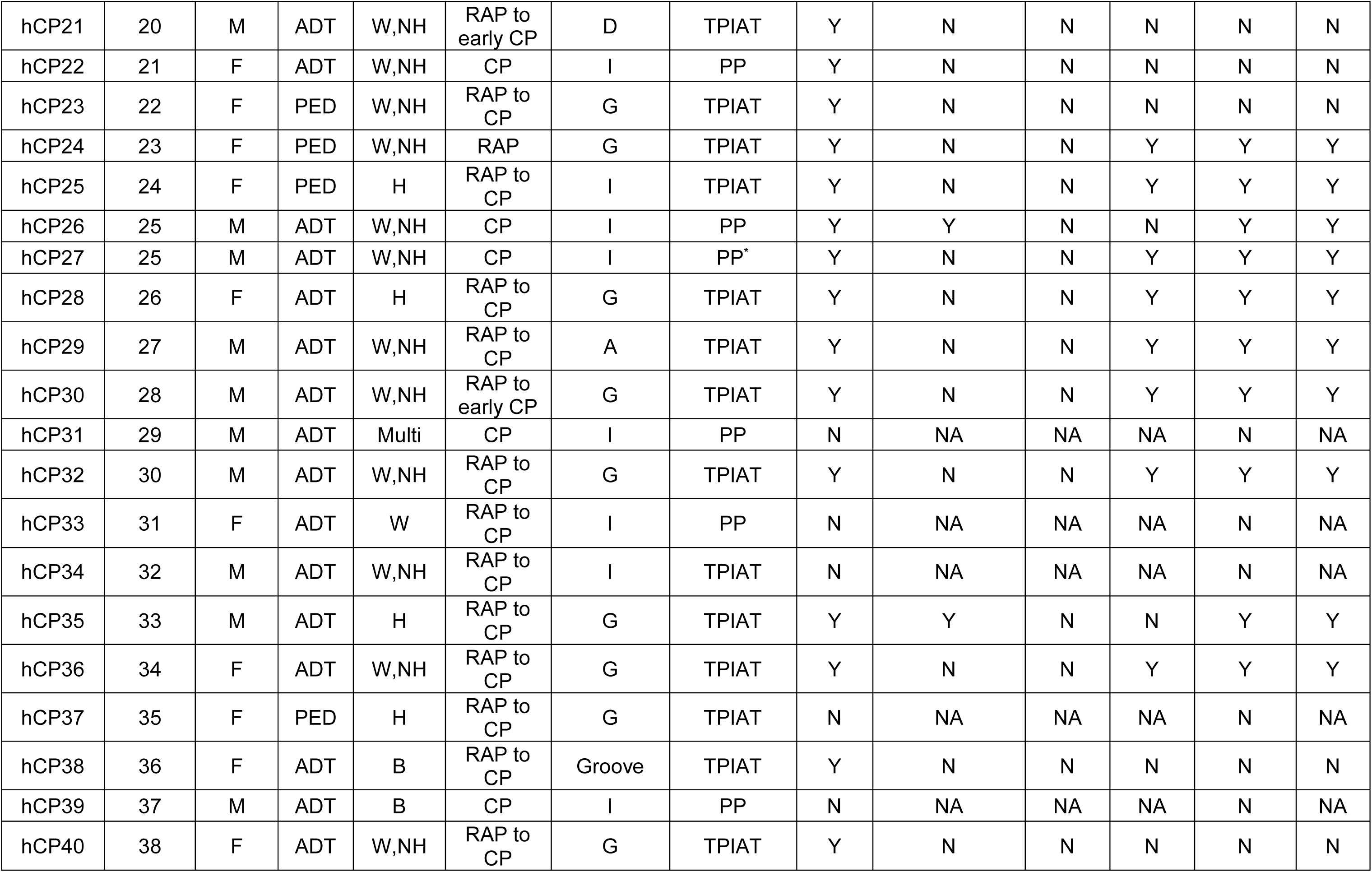
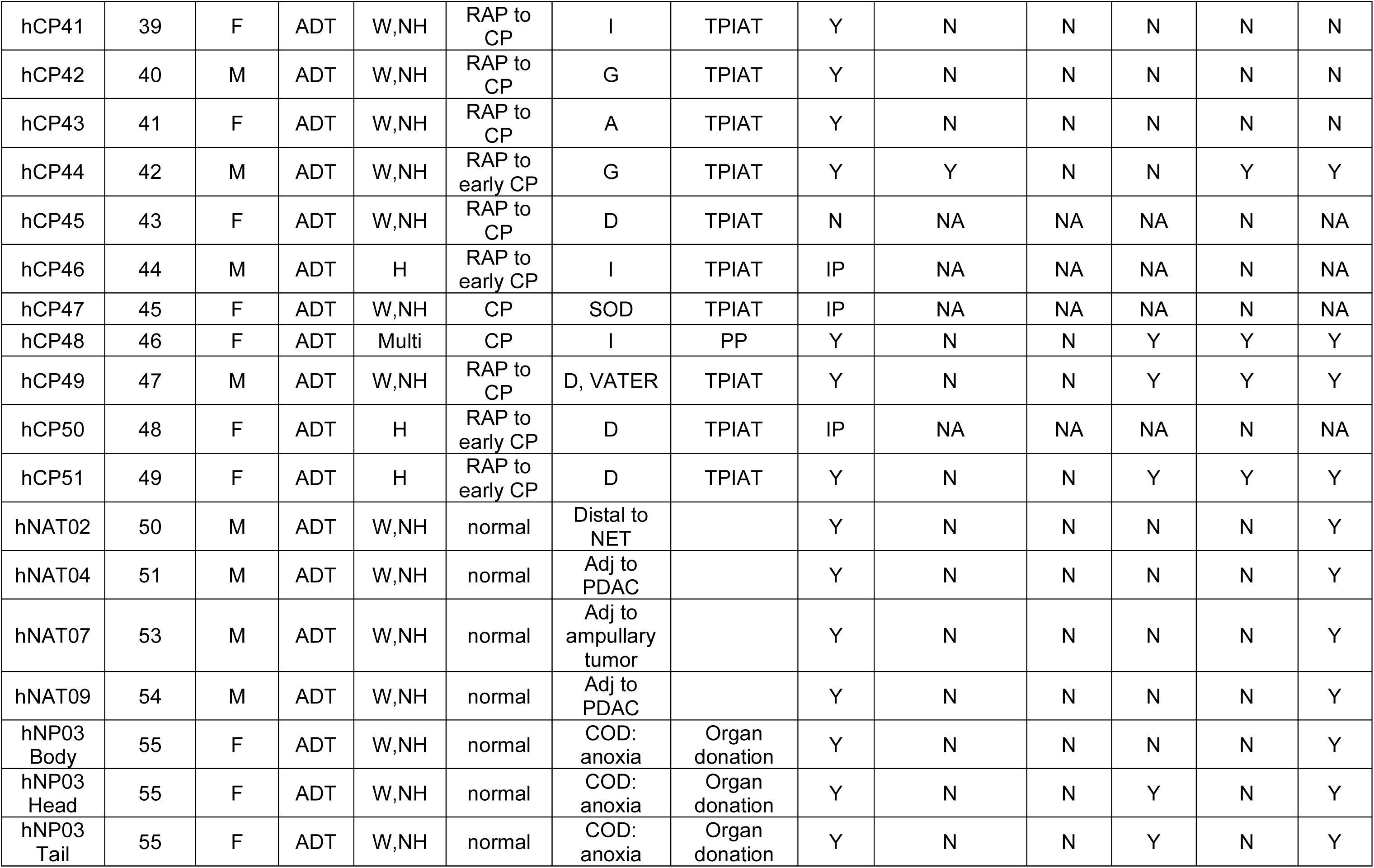
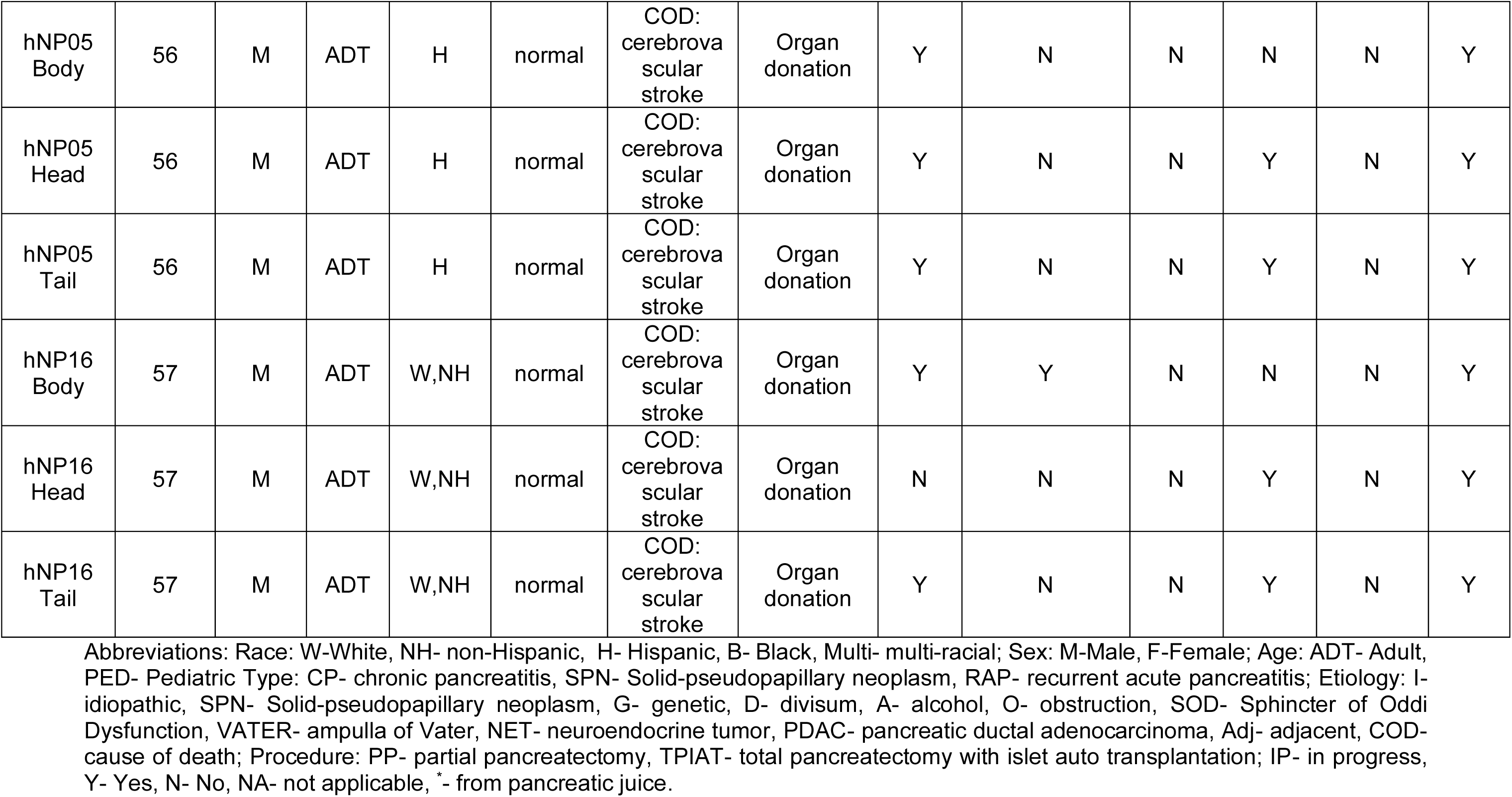
Patient sample information and downstream sequencing technologies applied to patient derived organoids (PDOs).

## Notes

### Competing Interest Statement

J.Gibson, P. Greer, and D. Whitcomb are employees a of Ariel Precision Medicine.

### Summary of Updates

Authors affiliations updated and list expanded to include additional experiments.

