## Supplemental Figures and Tables for "Chronic pancreatitis patient-derived organoids reveal new paths to precision therapeutics"

**Table S1. Patient sample information and downstream sequencing technologies applied to patient derived organoids (PDOs).**

| Sample | Patient | Sex | Age | Race | Type | Etiology | Procedure | PDO | Targeted sequencing | WES-PDO | WGS-PDO | WGS-PT | RNA seq |
|---|---|---|---|---|---|---|---|---|---|---|---|---|---|
| hCP01 | 1 | F | ADT | W,NH | CP | I | PP | Y | Y | N | N | Y | Y |
| hCP02 | 2 | F | ADT | W,NH | SPN | SPN | PP | Y | N | N | N | N | N |
| hCP03 | 3 | F | ADT | W,NH | CP | I | PP | Y | Y | N | N | Y | Y |
| hCP04 | 3 | F | ADT | W,NH | CP | I | PP | N | NA | NA | NA | N | NA |
| hCP05 | 4 | F | ADT | W,NH | CP | I | PP | N | NA | NA | NA | N | NA |
| hCP06 | 5 | F | ADT | W | RAP | G, D | TPIAT | N | NA | NA | NA | N | NA |
| hCP07 | 6 | F | ADT | W,NH | CP | I | PP* | Y | N | N | Y | Y | Y |
| hCP08 | 7 | F | PED | W,NH | RAP to CP | G | TPIAT | Y | Y | Y | N | Y | Y |
| hCP09 | 8 | F | ADT | W,NH | CP | I | PP | Y | N | N | N | N | N |
| hCP10 | 9 | M | ADT | W,NH | CP | I | PP | N | NA | NA | NA | N | NA |
| hCP11 | 10 | F | ADT | W,NH | RAP to CP | G | TPIAT | Y | Y | Y | N | Y | Y |
| hCP12 | 11 | F | PED | W,NH | RAP to CP | G | TPIAT | Y | N | N | N | N | N |
| hCP13 | 12 | F | ADT | W,NH | RAP to CP | G | TPIAT | Y | N | N | N | N | N |
| hCP14 | 13 | M | PED | W,NH | RAP to CP | G | TPIAT | N | NA | NA | NA | N | NA |
| hCP15 | 14 | M | ADT | W,NH | RAP to CP | A | TPIAT | Y | Y | Y | N | Y | Y |
| hCP16 | 15 | F | ADT | W,NH | RAP to CP | O | TPIAT | Y | Y | N | Y | Y | Y |
| hCP17 | 16 | F | PED | W,NH | RAP to CP | G | TPIAT | Y | Y | Y | N | Y | Y |
| hCP18 | 17 | F | ADT | W,NH | CP | I | TPIAT | Y | N | N | Y | Y | Y |
| hCP19 | 18 | F | ADT | Multi | CP | I | PP | N | NA | NA | NA | N | NA |
| hCP20 | 19 | F | PED | W,NH | RAP to CP | I | TPIAT | Y | Y | Y | N | Y | Y |

| | | | | | | | | | | | | | |
|---|---|---|---|---|---|---|---|---|---|---|---|---|---|
| hCP21 | 20 | M | ADT | W,NH | RAP to early CP | D | TPIAT | Y | N | N | N | N | N |
| hCP22 | 21 | F | ADT | W,NH | CP | I | PP | Y | N | N | N | N | N |
| hCP23 | 22 | F | PED | W,NH | RAP to CP | G | TPIAT | Y | N | N | N | N | N |
| hCP24 | 23 | F | PED | W,NH | RAP | G | TPIAT | Y | N | N | Y | Y | Y |
| hCP25 | 24 | F | PED | H | RAP to CP | I | TPIAT | Y | N | N | Y | Y | Y |
| hCP26 | 25 | M | ADT | W,NH | CP | I | PP | Y | Y | N | N | Y | Y |
| hCP27 | 25 | M | ADT | W,NH | CP | I | PP* | Y | N | N | Y | Y | Y |
| hCP28 | 26 | F | ADT | H | RAP to CP | G | TPIAT | Y | N | N | Y | Y | Y |
| hCP29 | 27 | M | ADT | W,NH | RAP to CP | A | TPIAT | Y | N | N | Y | Y | Y |
| hCP30 | 28 | M | ADT | W,NH | RAP to early CP | G | TPIAT | Y | N | N | Y | Y | Y |
| hCP31 | 29 | M | ADT | Multi | CP | I | PP | N | NA | NA | NA | N | NA |
| hCP32 | 30 | M | ADT | W,NH | RAP to CP | G | TPIAT | Y | N | N | Y | Y | Y |
| hCP33 | 31 | F | ADT | W | RAP to CP | I | PP | N | NA | NA | NA | N | NA |
| hCP34 | 32 | M | ADT | W,NH | RAP to CP | I | TPIAT | N | NA | NA | NA | N | NA |
| hCP35 | 33 | M | ADT | H | RAP to CP | G | TPIAT | Y | Y | N | N | Y | Y |
| hCP36 | 34 | F | ADT | W,NH | RAP to CP | G | TPIAT | Y | N | N | Y | Y | Y |
| hCP37 | 35 | F | PED | H | RAP to CP | G | TPIAT | N | NA | NA | NA | N | NA |
| hCP38 | 36 | F | ADT | B | RAP to CP | Groove | TPIAT | Y | N | N | N | N | N |
| hCP39 | 37 | M | ADT | B | CP | I | PP | N | NA | NA | NA | N | NA |
| hCP40 | 38 | F | ADT | W,NH | RAP to CP | G | TPIAT | Y | N | N | N | N | N |

| | | | | | | | | | | | | | |
|---|---|---|---|---|---|---|---|---|---|---|---|---|---|
| hCP41 | 39 | F | ADT | W,NH | RAP to CP | I | TPIAT | Y | N | N | N | N | N |
| hCP42 | 40 | M | ADT | W,NH | RAP to CP | G | TPIAT | Y | N | N | N | N | N |
| hCP43 | 41 | F | ADT | W,NH | RAP to CP | A | TPIAT | Y | N | N | N | N | N |
| hCP44 | 42 | M | ADT | W,NH | RAP to early CP | G | TPIAT | Y | Y | N | N | Y | Y |
| hCP45 | 43 | F | ADT | W,NH | RAP to CP | D | TPIAT | N | NA | NA | NA | N | NA |
| hCP46 | 44 | M | ADT | H | RAP to early CP | I | TPIAT | IP | NA | NA | NA | N | NA |
| hCP47 | 45 | F | ADT | W,NH | CP | SOD | TPIAT | IP | NA | NA | NA | N | NA |
| hCP48 | 46 | F | ADT | Multi | CP | I | PP | Y | N | N | Y | Y | Y |
| hCP49 | 47 | M | ADT | W,NH | RAP to CP | D, VATER | TPIAT | Y | N | N | Y | Y | Y |
| hCP50 | 48 | F | ADT | H | RAP to early CP | D | TPIAT | IP | NA | NA | NA | N | NA |
| hCP51 | 49 | F | ADT | H | RAP to early CP | D | TPIAT | Y | N | N | Y | Y | Y |
| hNAT02 | 50 | M | ADT | W,NH | normal | Distal to NET | | Y | N | N | N | N | Y |
| hNAT04 | 51 | M | ADT | W,NH | normal | Adj to PDAC | | Y | N | N | N | N | Y |
| hNAT07 | 53 | M | ADT | W,NH | normal | Adj to ampullary tumor | | Y | N | N | N | N | Y |
| hNAT09 | 54 | M | ADT | W,NH | normal | Adj to PDAC | | Y | N | N | N | N | Y |
| hNP03 Body | 55 | F | ADT | W,NH | normal | COD: anoxia | Organ donation | Y | N | N | N | N | Y |
| hNP03 Head | 55 | F | ADT | W,NH | normal | COD: anoxia | Organ donation | Y | N | N | Y | N | Y |
| hNP03 Tail | 55 | F | ADT | W,NH | normal | COD: anoxia | Organ donation | Y | N | N | Y | N | Y |

| | | | | | | | | | | | | | |
|---|---|---|---|---|---|---|---|---|---|---|---|---|---|
| hNP05 Body | 56 | M | ADT | H | normal | COD: cerebrovascular stroke | Organ donation | Y | N | N | N | N | Y |
| hNP05 Head | 56 | M | ADT | H | normal | COD: cerebrovascular stroke | Organ donation | Y | N | N | Y | N | Y |
| hNP05 Tail | 56 | M | ADT | H | normal | COD: cerebrovascular stroke | Organ donation | Y | N | N | Y | N | Y |
| hNP16 Body | 57 | M | ADT | W,NH | normal | COD: cerebrovascular stroke | Organ donation | Y | Y | N | N | N | Y |
| hNP16 Head | 57 | M | ADT | W,NH | normal | COD: cerebrovascular stroke | Organ donation | N | N | N | Y | N | Y |
| hNP16 Tail | 57 | M | ADT | W,NH | normal | COD: cerebrovascular stroke | Organ donation | Y | N | N | Y | N | Y |

Abbreviations: Race: W-White, NH- non-Hispanic, H- Hispanic, B- Black, Multi- multi-racial; Sex: M-Male, F-Female; Age: ADT- Adult, PED- Pediatric Type: CP- chronic pancreatitis, SPN- Solid-pseudopapillary neoplasm, RAP- recurrent acute pancreatitis; Etiology: I- idiopathic, SPN- Solid-pseudopapillary neoplasm, G- genetic, D- divisum, A- alcohol, O- obstruction, SOD- Sphincter of Oddi Dysfunction, VATER- ampulla of Vater, NET- neuroendocrine tumor, PDAC- pancreatic ductal adenocarcinoma, Adj- adjacent, COD- cause of death; Procedure: PP- partial pancreatectomy, TPIAT- total pancreatectomy with islet auto transplantation; IP- in progress, Y- Yes, N- No, NA- not applicable, *- from pancreatic juice.

Table S2. Patient cohort demographics, PDO generation, pathologist review and experimental methods completed per sample.

| Line | PATIENT # | Sex | Age | Race | Type | Etiology | Sample | PDO | % MT + | Fibrosis Score | Acinar Atrophy | Ductal Changes | Inflammation | Postmortem changes | Electro-scapel Artifacts | Pathologist Comments | WES- PDO | Targeted Sequening | WGS-PDO | RNAseq | WGS- PT | FIS deficient | Cytokine Analysis |
|---|---|---|---|---|---|---|---|---|---|---|---|---|---|---|---|---|---|---|---|---|---|---|---|
| hCP01 | 1 | F | ADT | W, NH | CP | I | Surgery | Y | 54.5% | moderate | moderate | moderate | mild | moderate | Y | Focal severe fibrosis | N | Y | N | Y | Y | N | Y |
| hCP02 | 2 | F | ADT | W, NH | SPN | SPN | Surgery | Y | 12.8% | moderade | mild | mild | mild | moderate | N | Hard to evaluate because of processing artifacts | N | N | N | N | N | NA | N |
| hCP03 | 3 | F | ADT | W, NH | CP | I | Surgery | Y | 14.7% | moderate | | | | | | | N | Y | N | Y | Y | N | Y |
| hCP04 | 3 | F | ADT | W, NH | CP | I | Surgery | N | 14.7% | moderate | mild | mild | mild | moderate | N | | NA | NA | NA | NA | N | NA | NA |
| hCP05 | 4 | F | ADT | W, NH | CP | I | Surgery | N (Frozen) | 94.6% | severe | severe | severe | severe | moderate | Y | Fibrotic tissue. There is almost no pancreatic tissue in this slide | NA | NA | NA | NA | N | NA | NA |
| hCP06 | 5 | F | ADT | W | RAP | G, D | TPIAT | N | 27.8% | severe | severe | severe | moderate | moderate | N | Areas compatible with acute pancreatitis | NA | NA | NA | NA | N | NA | NA |
| hCP07 | 6 | F | ADT | W, NH | CP | I | Juice | Y | 88.5% | severe | severe | severe | mild | moderate | Y | Few remaining acini and islets | N | N | Y | Y | Y | NA | N |
| hCP08 | 7 | F | PED | W, NH | RAP to CP | G | TPIAT | Y | 67.2% | severe | severe | severe | moderate | severe | N | | Y | Y | N | Y | Y | Y | Y |
| hCP09 | 8 | F | ADT | W, NH | CP | I | Surgery | Y | 51.7% | severe | severe | severe | severe | mild | N | Islets, numerous | N | N | N | N | N | NA | N |
| hCP10 | 9 | M | ADT | W, NH | CP | I | Surgery | N | 96.5% | severe | severe | severe | mild | mild | Y | Histological artifacts associated with electroscalpel | NA | NA | NA | NA | N | NA | NA |
| hCP11 | 10 | F | ADT | W, NH | RAP to CP | G | TPIAT | Y | 27.6% | severe | severe | severe | moderate | moderate | N | | Y | Y | N | Y | Y | Y | Y |
| hCP12 | 11 | F | PED | | RAP to CP | G | TPIAT | Y | 65.9% | severe | severe | severe | moderate | moderate | N | | N | N | N | N | N | NA | N |
| hCP13 | 12 | M | ADT | W, NH | RAP to CP | G | TPIAT | Y | 43.8% | severe | severe | severe | moderate | moderate | N | | N | N | N | N | N | NA | N |
| hCP14 | 13 | M | PED | W, NH | RAP to CP | G | TPIAT | N (Frozen) | 43.1% | severe | severe | severe | moderate | moderate | Y | Histological artifacts associated with electroscalpel | NA | NA | NA | NA | N | NA | NA |
| hCP15 | 14 | M | ADT | W, NH | RAP to CP | A | TPIAT | Y | 33.3% | severe | severe | severe | moderate | mild | N | | Y | Y | N | Y | Y | NA | N |
| hCP16 | 15 | F | ADT | | RAP to CP | O | TPIAT | Y | 67.5% | severe | severe | severe | moderate | moderate | N | | N | Y | Y | Y | Y | Y | Y |
| hCP17 | 16 | F | PED | W, NH | RAP to CP | G | TPIAT | Y | 70.6% | severe | severe | severe | moderate | moderate | Y | | Y | Y | N | Y | Y | NA | N |
| hCP18 | 17 | F | ADT | W, NH | CP | I | TPIAT | Y | 59.4% | severe | severe | severe | moderate | moderate | N | | N | N | Y | Y | Y | NA | N |
| hCP19 | 18 | F | ADT | Multi | CP | I | Surgery | N | 59.1% | IP | | | | | | | NA | NA | NA | NA | N | NA | NA |
| hCP20 | 19 | F | PED | W, NH | RAP to CP | I | TPIAT | Y | 33.9% | IP | | | | | | | Y | Y | N | Y | Y | NA | N |
| hCP21 | 20 | M | ADT | W, NH | RAP to early CP | D | TPIAT | Y | 29.2% | severe | severe | severe | severe | severe | N | Focal necrosis | N | N | N | N | N | NA | N |
| hCP22 | 21 | F | ADT | W, NH | CP | I | Surgery | Y | 56.6% | IP | | | | | | | N | N | N | N | N | NA | N |
| hCP23 | 22 | F | PED | W, NH | RAP to CP | G | TPIAT | Y | 34.0% | severe | severe | severe | severe | severe | N | | N | N | N | N | N | NA | N |
| hCP24 | 23 | F | PED | W, NH | RAP | G | TPIAT | Y | 36.0% | severe | severe | severe | severe | moderate | N | | N | N | Y | Y | Y | NA | N |
| hCP25 | 24 | F | PED | H | RAP to CP | I | TPIAT | Y | 22.4% | severe | severe | severe | severe | moderate | N | | N | N | Y | Y | Y | NA | N |
| hCP26 | 25 | M | ADT | W, NH | CP | I | Surgery | Y | 38.1% | moderate | mild | mild | mild | mild | Y | | N | Y | N | Y | Y | Y | Y |
| hCP27 | 25 | M | ADT | W, NH | CP | I | Juice | Y | Pancreas Juice | Pancreas Juice | | | | | | | N | N | Y | Y | Y | NA | Y |
| hCP28 | 26 | F | ADT | H | RAP to CP | G | TPIAT | Y | 11.6% | severe | severe | severe | severe | severe | N | | N | N | Y | Y | Y | NA | N |

| | | | | | | | | | | | | | | | | | | | | | | | |
|---|---|---|---|---|---|---|---|---|---|---|---|---|---|---|---|---|---|---|---|---|---|---|---|
| hCP29 | 27 | M | ADT | W, NH | RAP to CP | A | TPIAT | Y | 80.3% | severe | severe | severe | severe | moderate | N | | N | N | Y | Y | Y | NA | N |
| hCP30 | 28 | M | ADT | W, NH | RAP to early CP | G | TPIAT | Y | 55.9% | severe | severe | severe | severe | severe | N | | N | N | Y | Y | Y | NA | N |
| hCP31 | 29 | M | ADT | Multi | CP | I | Surgery | N | 11.5% | mild | mild | mild | mild | mild | N | Histological artifacts associated with electroscalpel | NA | NA | NA | NA | N | NA | N |
| hCP32 | 30 | M | ADT | W, NH | RAP to early CP | G | TPIAT | Y | 71.8% | severe | severe | severe | severe | severe | N | | N | N | Y | Y | Y | NA | N |
| hCP33 | 31 | F | ADT | W | CP | I | Surgery | N | 68.2% | severe | severe | severe | mild | moderate | Y | surviving islets | NA | NA | NA | NA | N | NA | NA |
| hCP34 | 32 | M | ADT | W, NH | RAP to CP | I | TPIAT | N | 4.9% | severe | severe | severe | severe | severe | N | | NA | NA | NA | NA | N | NA | NA |
| hCP35 | 33 | M | ADT | H | RAP to CP | G | TPIAT | Y | 39.6% | severe | severe | severe | severe | severe | N | | N | Y | N | Y | Y | Y | Y |
| hCP36 | 34 | F | ADT | W, NH | RAP to CP | G | TPIAT | Y | 89.7% | severe | severe | severe | severe | severe | N | | N | N | Y | Y | Y | NA | Y |
| hCP37 | 35 | F | PED | H | RAP to CP | G | TPIAT | N | 40.3% | severe | severe | severe | severe | severe | N | | NA | NA | NA | NA | N | NA | N |
| hCP38 | 36 | F | ADT | B | RAP to CP | I | TPIAT | Y | 51.3% | severe | severe | severe | severe | severe | N | | N | N | N | N | N | NA | N |
| hCP39 | 37 | M | ADT | B | CP | I | Surgery | N | 57.7% | severe | severe | severe | mild | | Y | | NA | NA | NA | NA | N | NA | N |
| hCP40 | 38 | F | ADT | W, NH | RAP to CP | G | TPIAT | Y | 84.3% | severe | severe | severe | severe | severe | N | | N | N | N | N | N | NA | N |
| hCP41 | 39 | F | ADT | W, NH | RAP to CP | I | TPIAT | Y | 29.5% | severe | severe | severe | severe | severe | N | | N | N | N | N | N | NA | N |
| hCP42 | 40 | M | ADT | W, NH | RAP to CP | G | TPIAT | Y | 66.4% | severe | severe | severe | severe | severe | N | | N | N | N | N | N | NA | N |
| hCP43 | 41 | F | ADT | W, NH | RAP to CP | A | TPIAT | Y | 84.1% | severe | severe | severe | severe | severe | N | | N | N | N | N | N | NA | Y |
| hCP44 | 42 | M | ADT | W, NH | RAP to early CP | G | TPIAT | Y | 70.6% | severe | severe | severe | severe | severe | N | | N | Y | N | Y | Y | Y | Y |
| hCP45 | 43 | F | ADT | W, NH | RAP to CP | D | TPIAT | N | Histological failure | Histological failure | | | | | | | NA | NA | NA | NA | N | NA | NA |
| hCP46 | 44 | M | ADT | H | RAP to early CP | I | TPIAT | IP | 66.1% | severe | severe | severe | severe | moderate | N | | NA | NA | NA | NA | N | NA | N |
| hCP47 | 45 | F | ADT | W, NH | CP | SOD | TPIAT | IP | 46.7% | severe | severe | severe | severe | severe | N | | NA | NA | NA | NA | N | NA | N |
| hCP48 | 46 | F | ADT | Multi | CP | I | Surgery | Y | 55.7% | moderate | moderate | moderate | mild | no | N | | N | N | Y | Y | Y | N | Y |
| hCP49 | 47 | M | ADT | W, NH | RAP to CP | G | TPIAT | Y | 85.0% | severe | severe | severe | severe | severe | N | | N | N | Y | Y | Y | N | Y |
| hCP50 | 48 | F | PED | H | RAP to early CP | D, VATER | TPIAT | IP | 12.9% | moderate | moderate | moderate | moderate | moderate | N | Areas compatible with acute pancreatitis | NA | NA | NA | NA | N | NA | N |
| hCP51 | 49 | F | ADT | H | RAP to early CP | D | TPIAT | Y | 10.5% | moderate | moderate | moderate | moderate | severe | N | | N | N | Y | Y | Y | NA | Y |

Key: Abbreviations: Race: W-White, NH- non-Hispanic, H- Hispanic, B- Black, Multi- multi-racial; Sex: M-Male, F-Female; Age: ADT- Adult, PED- Pediatric Type: CP- chronic pancreatitis, SPN- Solid-pseudopapillary neoplasm, RAP- recurrent acute pancreatitis; Etiology: I- idiopathic, SPN- Solid-pseudopapillary neoplasm, G- genetic, D- divisum, A- alcohol, O- obstruction, SOD- Sphincter of Oddi Dysfunction, VATER- ampulla of Vater, NET- neuroendocrine tumor, PDAC- pancreatic ductal adenocarcinoma, Adj- adjacent, COD- cause of death; Procedure: PP- partial pancreatectomy, TPIAT- total pancreatectomy with islet auto transplantation; IP- in progress, MT%- Masson's trichrome, Y- Yes, N- No, NA- not applicable, *- from pancreatic juice

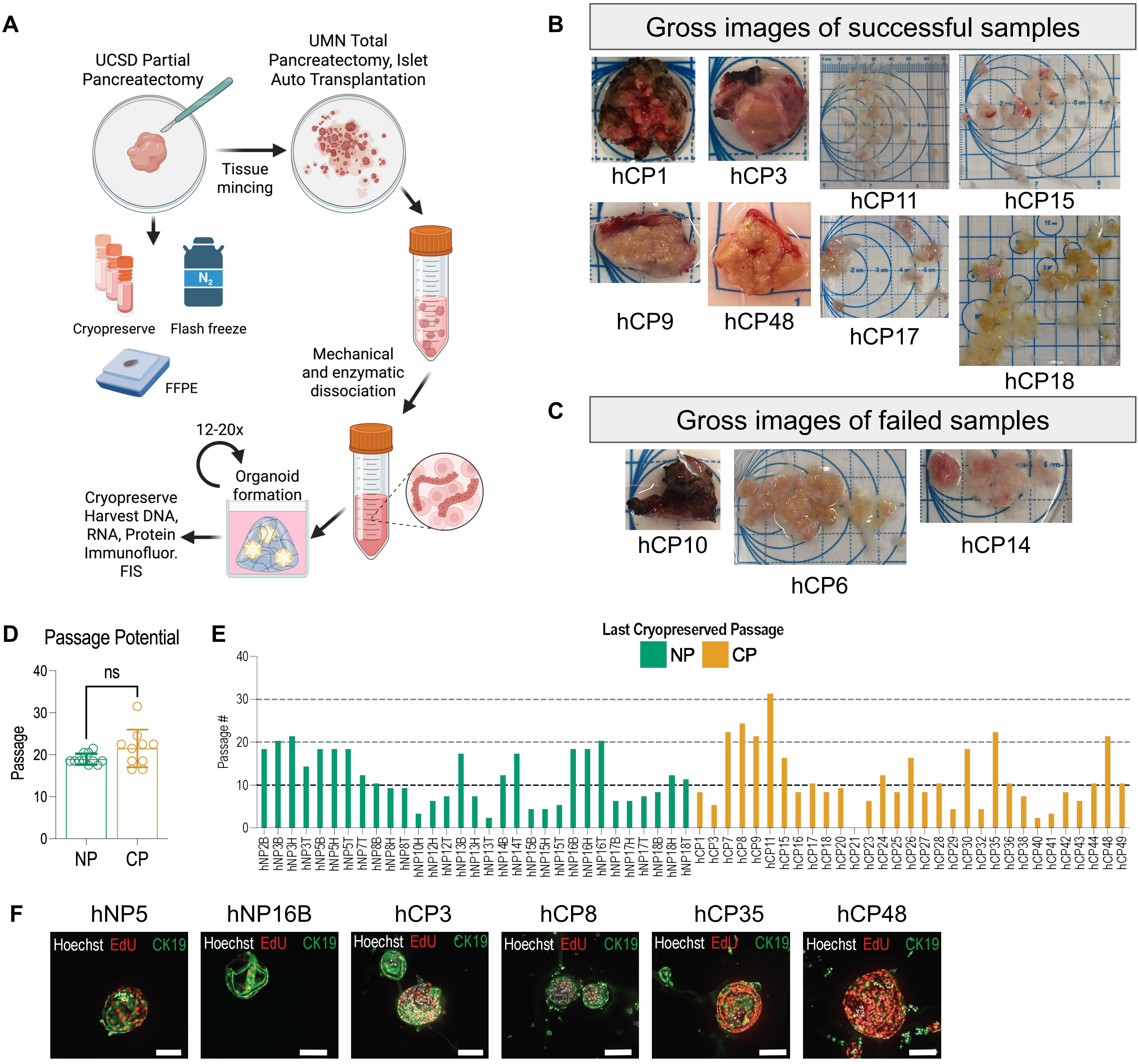


**Figure S1. Generation of organoids from human pancreas primary tissue and histological features of primary tissue that did not give rise to organoids, related to Figure 1.**
(A) Diagram of patient derived organoid generation from human pancreas specimen. Created in BioRender. Osorio Vasquez, V. (2025) https://BioRender.com/wsebx5k
(B) Gross images of CP samples that successfully produced organoids.
(C) Gross images of CP samples that failed to produce organoids.
(D) Number of days that passed between passaging and expanding either NP or CP organoids are not different. P-value determined by Mann-Whitney test.
(E) Last cryopreserved passage for NP and CP organoids.
(F) Whole mount immunofluorescence of PDOs stained with Hoechst (nuclei), EdU (pulsed for 25 hours), and CK19 (ductal marker). Scale bar 100 µm. Quantified in Figure 1G.

A

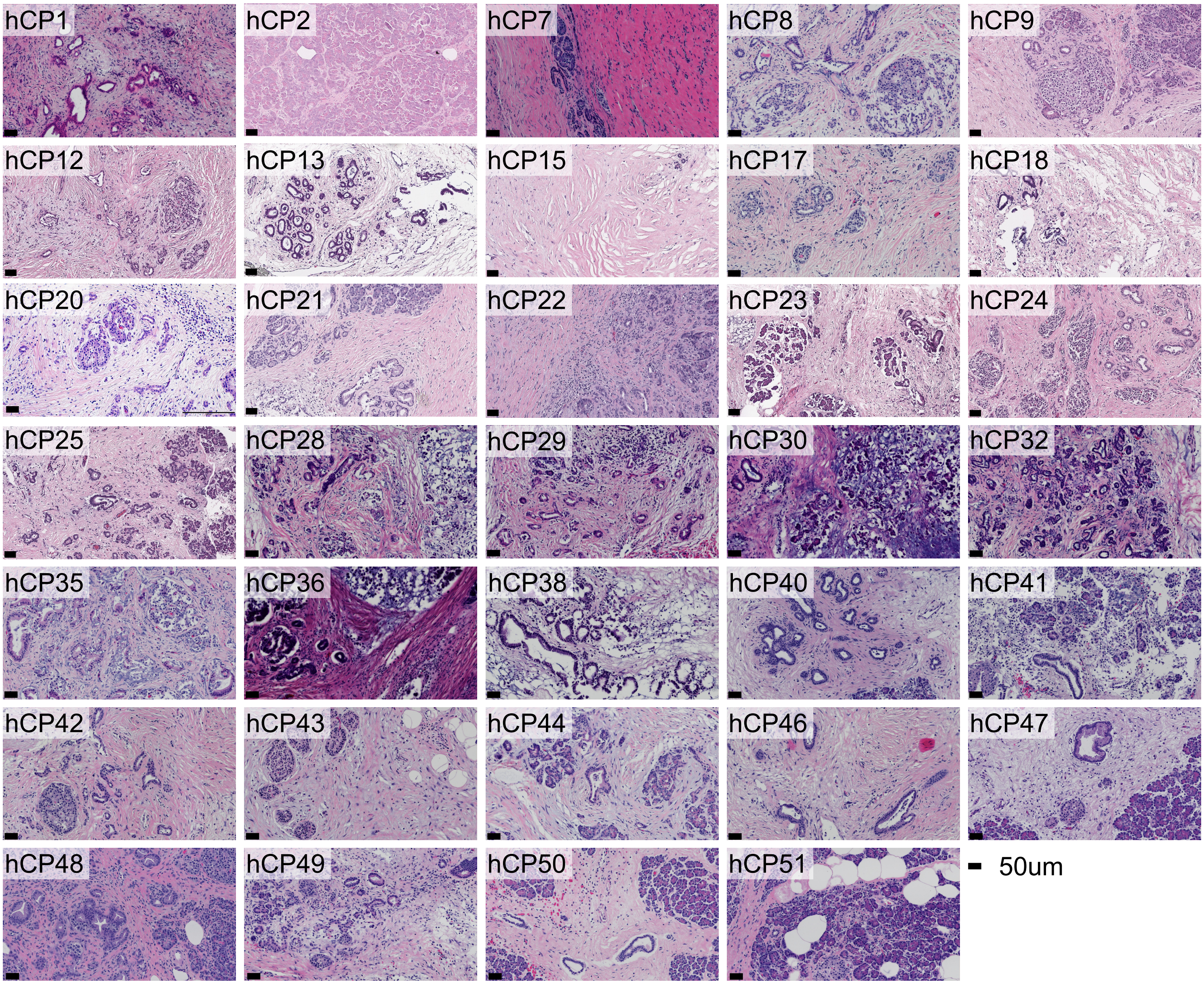


B

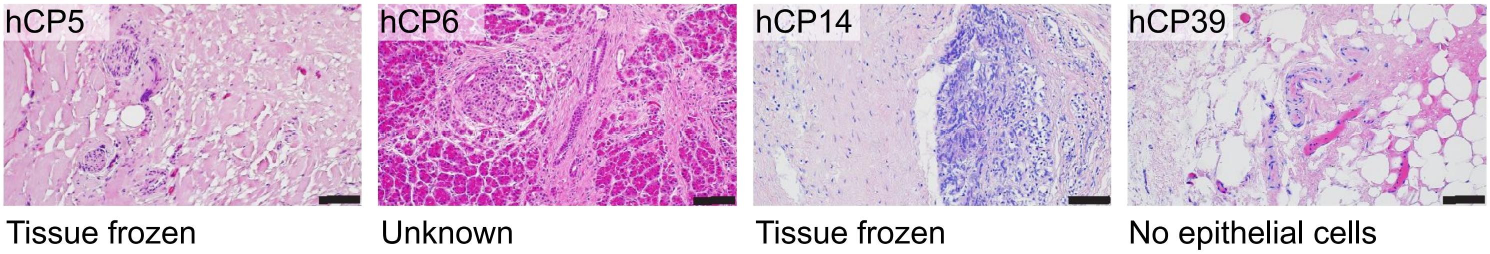


**Figure S2. Generation of organoids from human pancreas primary tissue and histological features of primary tissue that did not give rise to organoids, related to Figure 1.**

(A) H&E of primary specimens that formed organoids. Scale bar 50 µm.

(B) H&E of primary specimens that did not form organoids. Lack of epithelial cells, high degrees of fibrosis, fat cell replacement and samples that arrived frozen are represented. Scale bar 100 µm.

A

| hCP Pair | Match Rate (%) | ICC | 95% CI (±) |
|---|---|---|---|
| hCP07 | 98.34 | 0.9644 | 0.0001 |
| hCP08 | 98.33 | 0.8335 | 0.0261 |
| hCP11# | 90.77 | 0.8119 | 0.0195 |
| hCP15# | 91.59 | 0.8286 | 0.0264 |
| hCP16# | 92.17 | 0.8415 | 0.0291 |
| hCP17# | 91.79 | 0.8326 | 0.0252 |
| hCP18 | 98.33 | 0.9640 | 0.0001 |
| hCP20# | 90.98 | 0.8166 | 0.0223 |
| hCP24 | 98.44 | 0.9663 | 0.0001 |
| hCP25 | 98.49 | 0.9678 | 0.0000 |
| hCP27 | 98.45 | 0.9664 | 0.0000 |
| hCP28 | 98.43 | 0.9665 | 0.0000 |
| hCP29 | 98.42 | 0.9659 | 0.0001 |
| hCP30 | 98.46 | 0.9670 | 0.0000 |
| hCP32 | 98.43 | 0.9659 | 0.0001 |
| hCP36 | 98.40 | 0.9656 | 0.0000 |
| hCP48 | 98.40 | 0.9659 | 0.0001 |
| hCP49 | 98.39 | 0.9657 | 0.0000 |
| hCP51 | 98.44 | 0.9665 | 0.0001 |
| hCP30 vs hCP48 | 71.85 | 0.3986 | 0.0007 |
| Primary Tissue 17 vs hCP32 | 73.95 | 0.4344 | 0.0008 |
| hCP7 vs hCP20# | 56.62 | 0.1148 | 0.0457 |

B

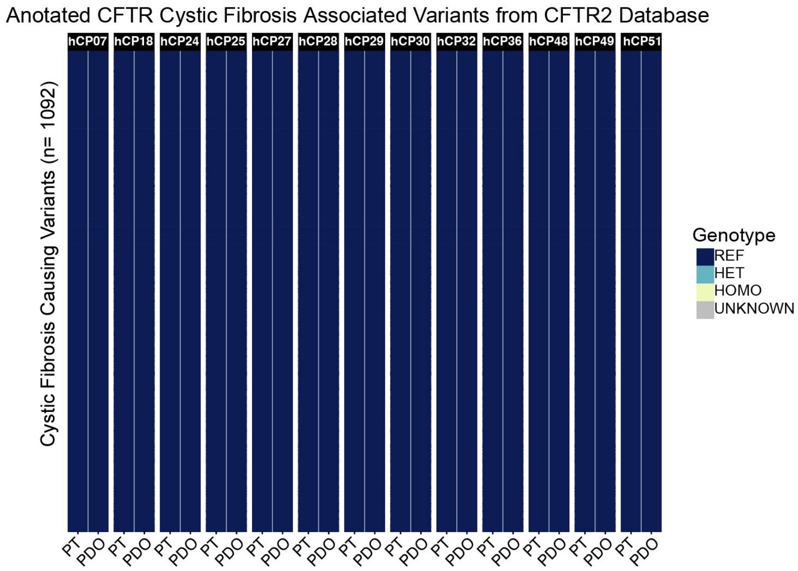


C

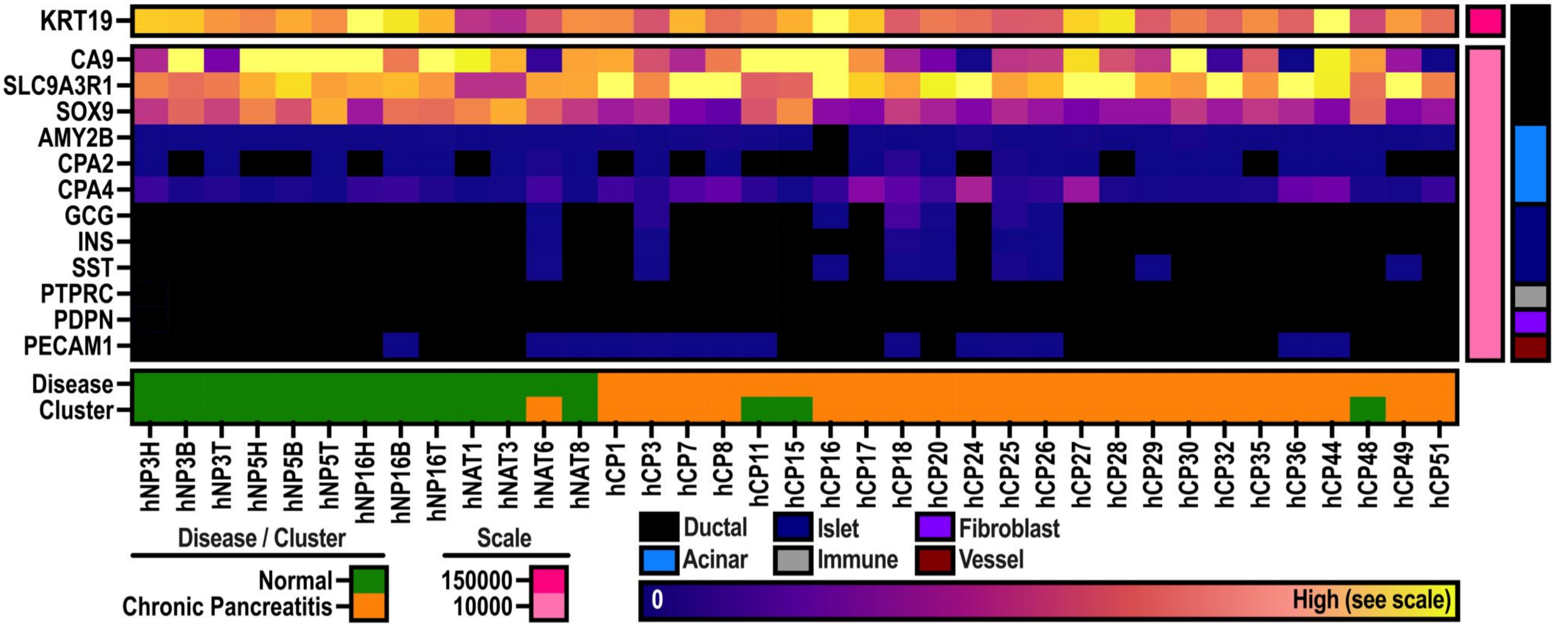


**Figure S3. CP PDOs genetic concordance, lack of cystic fibrosis associated variants of CFTR, related to Figure 2, and hCP PDO lineage inferred from transcriptional data; related to Figure 3.**

(A) Genetic concordance, intraclass correlation coefficient (ICC), and 95% confidence interval determined from single nucleotide polymorphisms (SNPs) of PDO and matched primary tissue specimens by whole genome sequencing (WGS) or whole exome sequencing (#). For additional patient sample sequencing modality information see also Table S1.
(B) Cystic fibrosis causing variants annotated in CFTR2 database (n=1092) are not expressed in CP organoids.
(C) Normalized counts from RNAseq for ductal, acinar, endocrine, fibroblast, endothelial, and immune cell lineage markers. Clusters and specimen disease/origin are annotated.

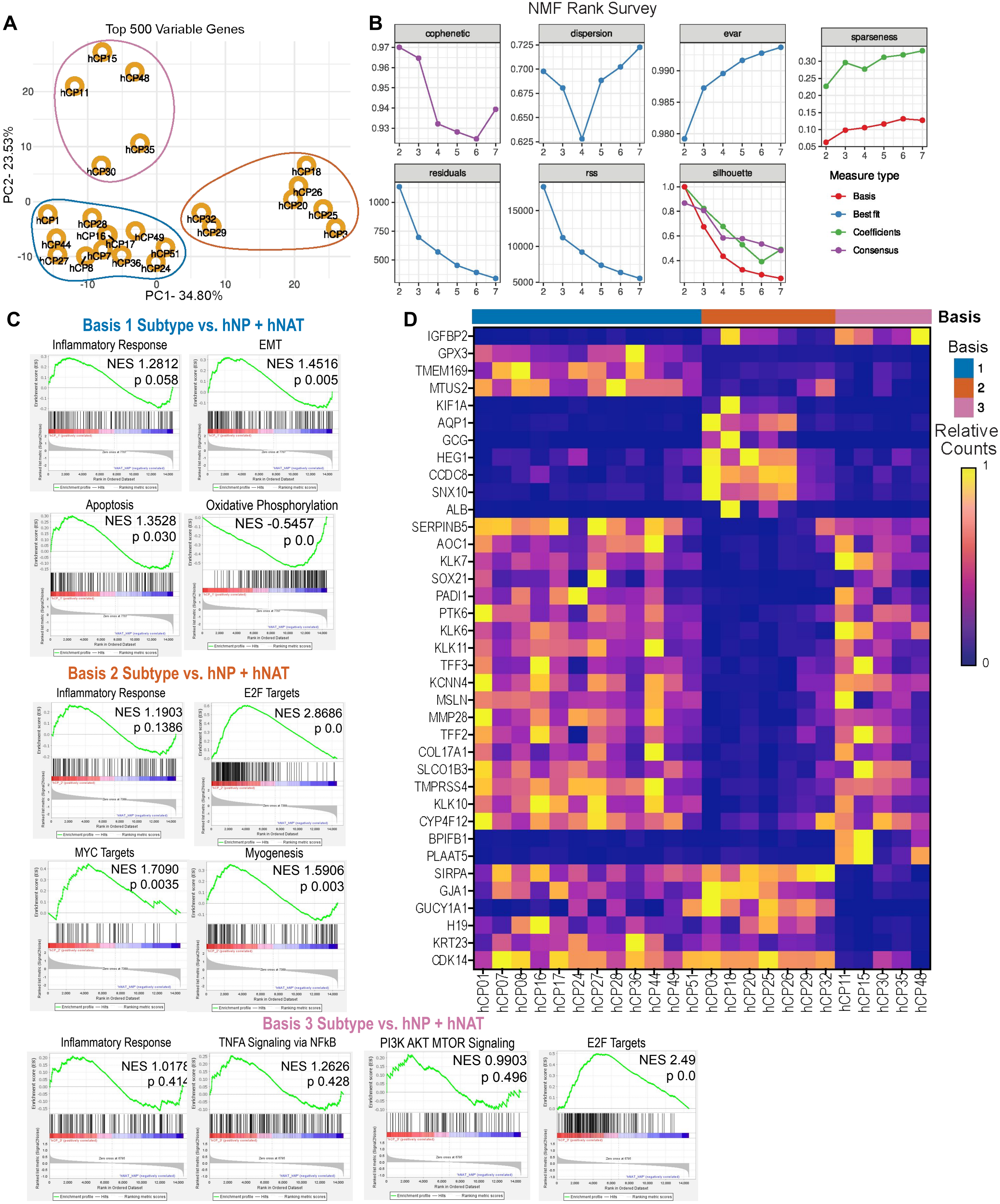


**Figure S4. Transcriptionally distinct subtypes of CP generated from non-negative matrix factorization (NMF) analysis, related to Figure 3.**

(A) Principal component analysis of transcriptomic data from CP organoids. Subtypes defined by basis are shown.

(B) Cophenetic, dispersion, evar, residuals, rss, silhouette, and spareness detailed from NMF rank survey. Basis 3 was selected as encapsulating the data best.

(C) Gene set enrichment analysis (GSEA) of each CP subtype compared to NP and hNAT PDOs. Normalized enrichment score (NES) and nominal p-value (nom. p-value) annotated.

(D) Differentially expressed genes (p value <0.05, |log2fold change| > 1) between three subtypes of hCP with Kim-scores greater than or equal 0.07 for basis 1 and 2, and greater than or equal to 0.06 for basis 3. Relative variance stabilized counts plotted.

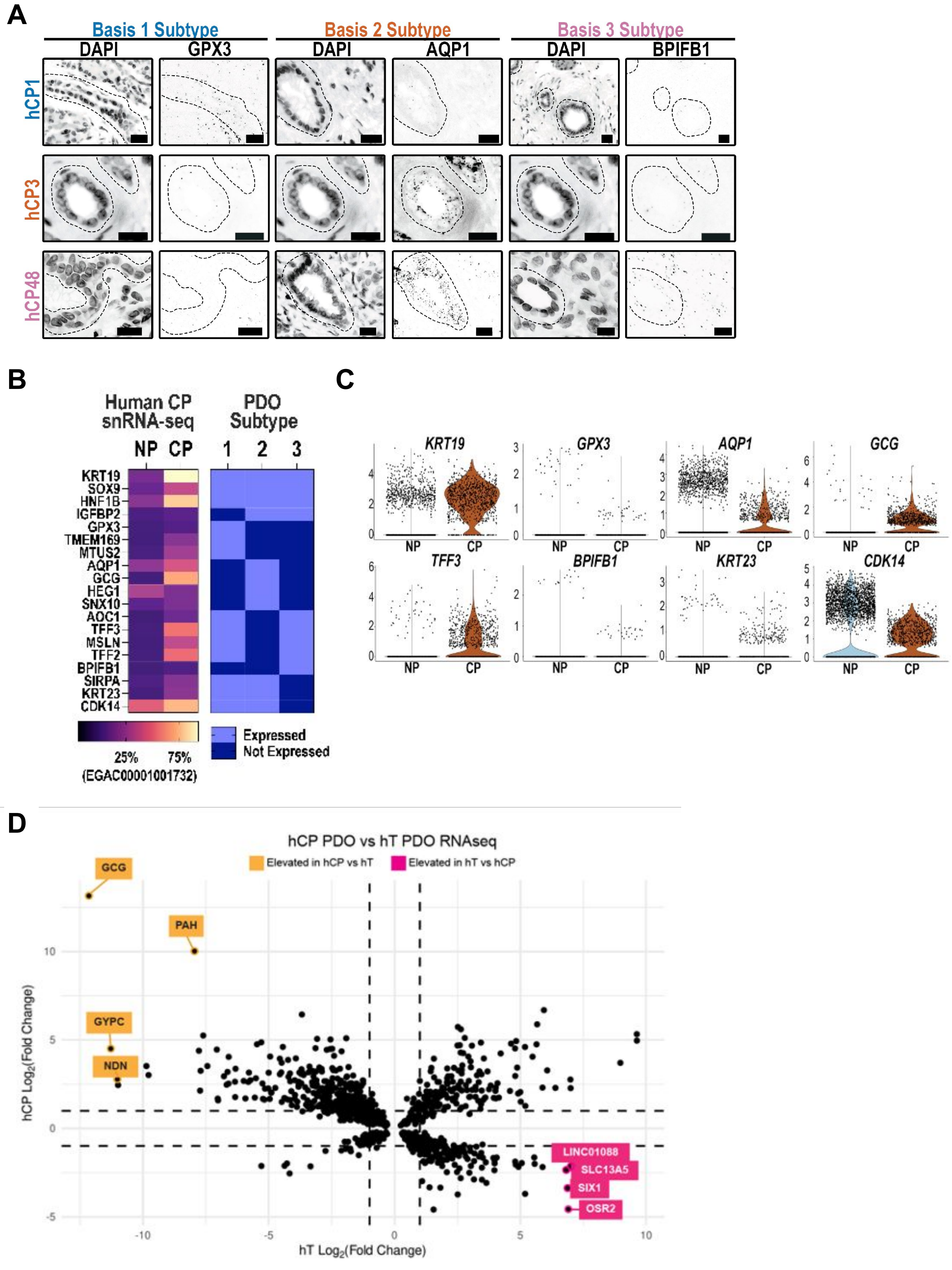


**Figure S5. CP subtype validation, and differences compared to human pancreatic cancer organoids, related to Figure 3.**

(A) Fluorescent RNA in situ hybridization of primary tissue sections of CP. Ducts are annotated in dashed lines in both DAPI and gene probe images. Scale bar 20 µm. Quantitation of probe hybridization per nuclei for each field of view. For expression of subtype defining genes in an independent cohort see Figure S5C.

(B) Percentage of cells from the normal adjacent to tumor (NP) or CP ductal cell cluster expressing the ductal marker KRT19 and genes driving the transcriptional subtypes from snRNA-seq data (EGAC00001001732). A key for CP PDO subtype gene associations from Figure 4 represented as expressed or not expressed.

(C) Normalized counts (nCounts) of subtype defining genes and ductal expression markers in NP and CP patient samples from EGAC00001001732 data. Each dot is a single nucleus.

(D) Cloverleaf plot of differentially expressed genes comparing CP organoids and pancreatic cancer organoids to NP organoids. Dashed lines signify $log_2$Fold change =1. Annotated genes are enriched in PDA or CP organoids. hT: PDA organoids.

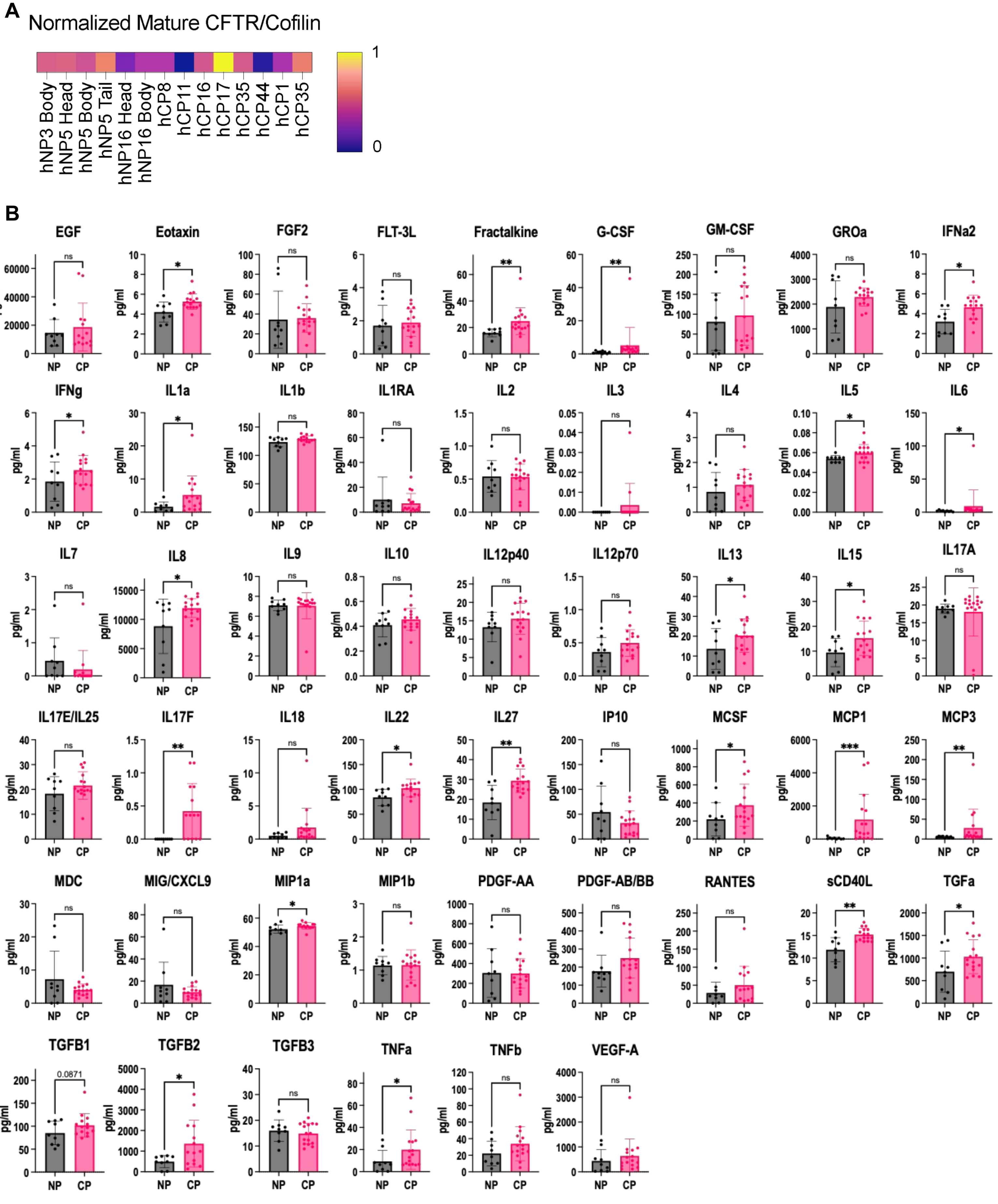


**Figure S6. CFTR expression in hNP organoids and secreted proteins assayed in NP and CP samples, related to Figure 4.**

(A) Expression of mature CFTR in hNP and hCP organoids quantified relative to cofilin loading control.

(B) Secreted proteins quantified in conditioned media from NP or CP organoids. P-value determined by Mann-Whitney test.

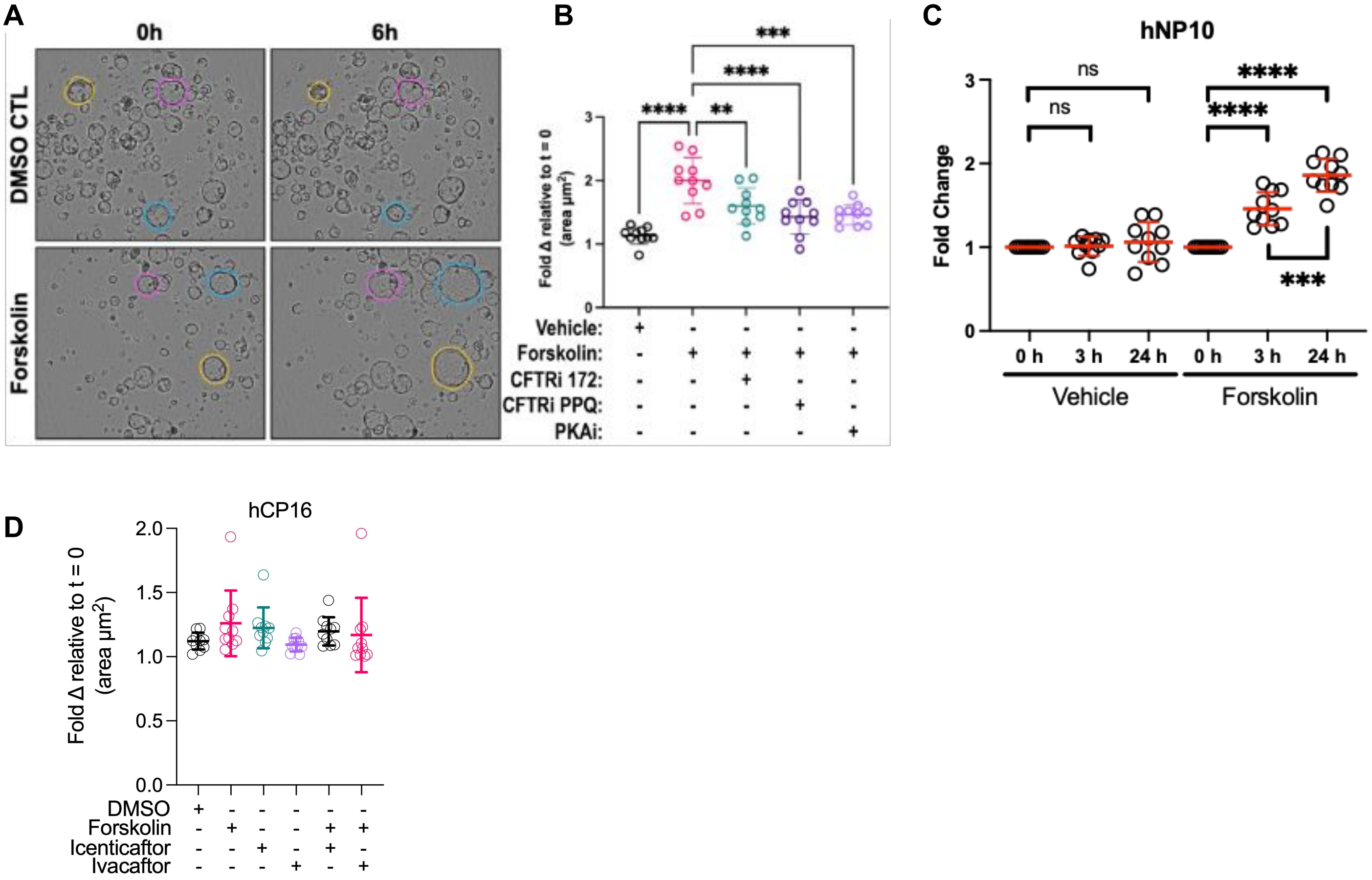


**Figure S7. NP organoids are proficient in CFTR dependent swelling and some CP organoids genotypes are swelling deficient even upon addition of CFTR potentiators, related to Figure 5.**

(A) Representative brightfield images of mouse normal organoids treated with forskolin for 6 hours.

(B) Mouse normal (mNP11) organoids treated with forskolin for 6 hours and pre-treated with CFTR inhibitors (CFTRi-172, PPQ-102), or protein kinase A inhibitor (H-89) for 2 hours. Swelling is inhibited with the addition of CFTR and PKA inhibitors.

(C) Human normal (NP) organoids treated with forskolin swell overtime. Represented as fold change in area of organoids.

(D) hCP16 organoids do not swell in response to 6 hour treatment with forskolin or CFTR potentiators.
